# An NF-κB/IRF1 axis programs cDC1s to drive anti-tumor immunity

**DOI:** 10.1101/2020.12.28.424621

**Authors:** G Ghislat, AS Cheema, E Baudoin, C Verthuy, PJ Ballester, K Crozat, N Attaf, C Dong, P Milpied, B Malissen, N Auphan-Anezin, TP Vu Manh, M Dalod, T Lawrence

## Abstract

Conventional type 1 dendritic cells (cDC1s) are critical for anti-tumor immunity. They acquire antigens from dying tumor cells and cross-present them to CD8^+^ T cells, promoting the expansion of tumor-specific cytotoxic T cells. However, the signaling pathways that govern the anti-tumor functions of cDC1s are poorly understood. We mapped the molecular pathways regulating intra-tumoral cDC1 maturation using single cell RNA sequencing. We identified NF-κB and IFN pathways as being highly enriched in a subset of functionally mature cDC1s. The specific targeting of NF-κB or IFN pathways in cDC1s prevented the recruitment and activation of CD8^+^ T cells and the control of tumor growth. We identified an NF-κB-dependent IFNγ-regulated gene network in cDC1s, including cytokines and chemokines specialized in the recruitment and activation of cytotoxic T cells. We used single cell transcriptomes to map the trajectory of intra-tumoral cDC1 maturation which revealed the dynamic reprogramming of tumor-infiltrating cDC1s by NF-κB and IFN signaling pathways. This maturation process was perturbed by specific inactivation of either NF-κB or IRF1 in cDC1s, resulting in impaired expression of IFN-γ-responsive genes and consequently a failure to efficiently recruit and activate anti-tumoral CD8^+^ T cells. Finally, we demonstrate the relevance of these findings to cancer patients, showing that activation of the NF-κB/IRF1 axis in association with cDC1s is linked with improved clinical outcome. The NF-κB/IRF1 axis in cDC1s may therefore represent an important focal point for the development new diagnostic and therapeutic approaches to improve cancer immunotherapy.

**One Sentence Summary:** NF-κB and IRF1 coordinate intra-tumoral cDC1 maturation and control of immunogenic tumor growth.

## Introduction

Immunotherapy has brought unprecedented advances in cancer treatment, yet still the majority of patients don’t respond to current therapies. Hence, there is a desperate need to further understand what dictates patient responses to immunotherapy and design new approaches to improve therapeutic outcomes.

Previous studies in several preclinical cancer models support an important contribution of conventional type 1 dendritic cells (cDC1s) to effective immunotherapy, including adoptive T cell therapy and immune-checkpoint blockade (*1–3*). Furthermore, the frequency of cDC1s in human cancers is associated with improved prognosis and response to immunotherapy (*4*). Importantly, cDC1s in humans and mice share development pathways and phenotypic markers, including expression of the chemokine receptor XCR1 as a lineage marker (*4, 5*). The defining property of cDC1s is the cross-presentation of antigens via the class I (major histocompatibility complex) MHC pathway to CD8^+^ T cells (*6*) which has been shown to be critical for anti-tumor immune responses (*7*). In line with this function, cDC1s express a number of molecules associated with the uptake and processing of antigens from dying cells for cross-presentation (*8*). But cross-presentation alone is not sufficient for the cDC1-dependent control of tumor growth (*9*), implying other functions of cDC1s play important roles in anti-tumor immunity. For example, DCs can be important sources of chemokines and cytokines that promote CD8^+^ T cell recruitment and activation in tumors. However, the tumor-microenvironment suppresses the recruitment and activation of cDC1s, through various mechanisms, which contributes to evasion of anti-tumor immunity and may underly resistance to immunotherapy in cancer patients (*4*). The intrinsic molecular pathways regulating intra-tumoral cDC1 functions remain poorly understood but are critical to determine how cDC1s can be targeted to improve cancer immunotherapy.

Previous studies have been hampered by the lack of experimental tools to dissect the specific functions of cDC1s, mostly relying on the use of Batf3-deficient mice that lack cDC1s, which precludes the study of pathways regulating cDC1 functions within tumors. The recent development of transgenic mice allowing the specific targeting of cDC1s, based on their unique expression of the chemokine receptor XCR1 (*4, 5*), has revealed important new aspects of cDC1 functions in tumors. For example, in a recent study the targeted deletion of MHC II in Xcr1-expressing cells revealed an unexpected role in priming CD4^+^ T cells, which was required for rejection of transplanted tumors bearing a xenoantigen (*10*).

The roles of cDC1s in anti-tumor immunity have mostly been addressed in the context of immunotherapy. Several reports have described the importance of cDC1s in various preclinical models for effective immunotherapy, including adoptive T cell therapy (*1,2*) and immune checkpoint blockade (*3,11–15*). In the context of adoptive T cell therapy, tumor-resident cDC1s were reported to produce the CXCR3-dependent chemokines CXCL9 and CXCL10, required for the recruitment of effector CD8^+^ T cells (*1*). In another study, increased IFN-γ production during anti-PD-1 therapy, was shown to drive IL-12 expression by DCs leading to increased CD8^+^ T cell activation (*16*). These studies demonstrated the important contribution of cDC1s to anti-tumor immune responses in the presence of activated CD8^+^ T cells producing IFNγ.

Tumor immunogenicity and the recruitment of tumor-infiltrating CD8^+^ T cells is a critical factor for clinical responses to immunotherapy (*17–19*). Tumor antigens broadly fall into two categories; those specifically expressed by cancer cells, including neoantigens generated by genetic alterations, and those that are highly but not exclusively expressed by cancer cells. Neoantigens, associated with cancers with high mutational burden, exert the strongest T cell-mediated anti-tumor immunity, which often correlates with improved responses to immunotherapy (*20*). However, the roles of cDC1s in relation to intrinsic tumor-immunogenicity have not yet been addressed.

We investigated the molecular pathways in tumor-infiltrating cDC1s that govern their anti-tumor functions in relation to tumor-immunogenicity. Using genetic tools to specifically target cDC1s, we demonstrate their specific role in the control of immunogenic tumor growth. Single-cell RNA sequencing revealed the trajectory of intra-tumoral cDC1 maturation and identified distinct activation states associated with tumor-immunogenicity. By targeting specific signaling pathways in cDC1s, we revealed molecular pathways that drive intra-tumoral cDC1 maturation leading to the recruitment and activation of anti-tumoral CD8^+^ T cells. Finally, we show that activation of these pathways in human cancer strongly correlates with good prognosis. Therefore, manipulating these pathways could offer new therapeutic strategies to overcome resistance to immunotherapy and improve outcomes for cancer treatment.

## Results

### cDC1s are required for control of immunogenic tumors

To explore the role of cDC1s in relation to tumor-immunogenicity, we used a clinically relevant mouse melanoma model based on expression of Braf^V600E^ and deletion of tumor suppressors *Pten* and *Cdkn2a,* specifically in melanocytes (YUMM1.7) (*21, 22*). Previous studies have established that YUMM1.7 tumors are not immunogenic and do not show increased growth in Rag-deficient mice, which lack T and B lymphocytes (*18*). Consequently, these tumors are also resistant to immune-checkpoint therapy. We exploited the fact YUMM1.7 tumors derived from male mice express the minor histocompatibility antigen HY (*23*), to assess tumor growth in an allogeneic setting by transplanting YUMM1.7 tumors in syngeneic female mice. Although tumors progressed in both male and female mice, tumor growth rate was significantly reduced in females (Fig. 1A), which was associated with a dramatic increase in leukocyte infiltration (Fig.S1A), suggesting that YUMM1.7 tumors were immunogenic in female mice. Indeed, analysis of tumor-infiltrating CD8^+^ T cells showed a substantial increase in recruitment of effector CD8^+^ T cells expressing IFN-γ and TNF-α in tumors from female mice compared to males (Fig. 1B). In addition, analysis of gene expression in tumorinfiltrating CD8^+^ T cells from female mice revealed increased expression of genes underpinning effector functions (*21, 22*) (Fig.S1B). Analysis of cytokines in tumor lysates also showed an increased production of various chemokines and inflammatory cytokines in tumors from female mice compared to males (Fig.S1C,D). To confirm the specificity of the T cell response in female mice, we next assessed the frequency of CD8^+^ T cells recognizing the tumor-associated HY antigen, by using MHC-I tetramers. As expected, HY-specific CD8^+^ T cells were absent from YUMM1.7 tumors in male mice, but were readily detected in tumors from females (Fig.1C). These data clearly established the intrinsic immunogenicity of YUMM1.7 tumors in female mice.

**Figure 1.**
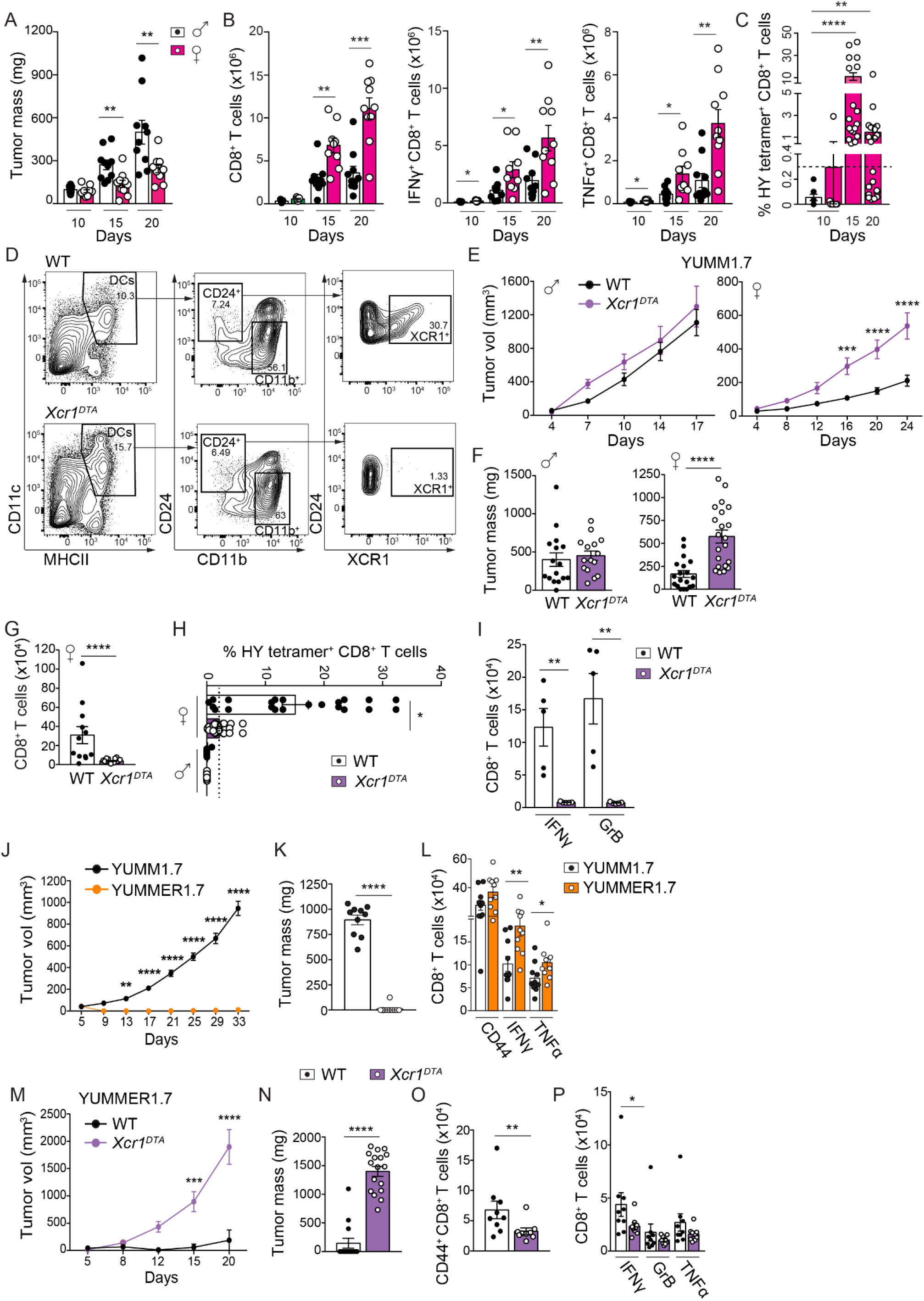
cDC1s are required for control of immunogenic tumor growth. (**A**) YUMM1.7 tumor growth in male and female mice. (**B**) Quantification of tumor-infiltrating CD8^+^ T cells, and IFN-γ or TNF-α producing cells by flow cytometry in YUMM1.7 tumors from male and female mice. (**C**) Tetramer analysis of HY-specific CD8^+^ T cells (Smcy_(738-746)_/H-2Db and Uty_(243-254)_/H-2Db) in YUMM1.7 tumors. The dashed line represents the highest frequency obtained with an irrelevant control epitope (HPV16 E7(49-57)/H-2Db). (**D**) Representative flow cytometry analysis of cDC1 in YUMM1.7 tumors (gated as CD3^-^ CD19^-^ NK1.1^-^ Ly6G^-^ CD11c^+^ MHCII^+^ CD64^-^) from female WT and *Xcr1^DTA^* mice. (**E**,**F**) YUMM1.7 tumor volume (**E**) and mass (**F**) in male and female WT or *Xcr1^DTA^* mice. In (**E**), n=5; 5; 11; 10 in male WT; *Xcr1^DTA^;* female WT; *Xcr1^DTA^* mice respectively. (**G-I**) Quantification of total CD8^+^ T cells (**G**), frequency of HY-tetramer positive CD8^+^ T cells (**H**) and IFN-γ or Granzyme B (Grb) producing cells (**I**) in YUMM1.7 tumors from WT and *Xcr1^DTA^* mice. (**J,K**) Growth of YUMM1.7 and YUMMER1.7 tumors in female and male mice, respectively. In (**J**), n=10 in YUMM1.7 and YUMMER1.7, respectively. (**L**) Quantification of CD44^+^ CD8^+^ T cells and IFN-γ or TNF-α producing cells in tumor-draining lymph nodes (TDLN) from mice engrafted with YUMM1.7 and YUMMER1.7 tumors. (**M,N**) YUMMER1.7 tumor volume (**M**) and mass (**N**) in male WT or *Xcr1^DTA^* mice. In (**M**), n=9 in WT and*Xcr1^DTA^* mice. (**O**,**P**) Quantification of CD44^+^ CD8^+^ T cells (**O**) and IFN-γ, Grb or TNF -α producing cells (**P**) in TDLN from mice engrafted with YUMMER1.7 tumors. Total cell numbers are indicated for 250mg of tumor tissue. Pooled data from three independent experiments or representative data from at least two independent experiments are shown. Data are represented as mean ± SEM. In scatter plots each point corresponds to an individual mouse. Statistical analysis was performed by two-way ANOVA followed by Sidak’s multiple comparison tests (**E, J** and **M**) or Mann-Whitney test. Fisher’s exact test was used for frequency of tetramer positive cells when their proportion > the highest level obtained with HPV16 E7_(49-57)_/H-2Db tetramer (**C** and **H**).

Taking advantage of the difference in immunogenicity between YUMM1.7 tumors in male and female mice, we proceeded to evaluate the specific contribution of cDC1s to the control of tumor growth. Previous studies implicating cDC1s in anti-tumor immune responses and immunotherapy in preclinical models had used *βatf3*-deficient mice constitutively lacking cDC1s. However, Batf3 is not exclusively expressed by cDC1s and these mice display other phenotypes in addition to the absence cDC1s, including intrinsic defects in memory CD8^+^ T cells (*24, 25*), reduced numbers of cDC2s (*14*) and an increased frequency of regulatory T cells (Treg) (*26*). In contrast, the chemokine receptor Xcr1 is a specific marker for cDC1s conserved in humans and mice (*4, 5*). Thus, to examine the specific role of cDC1s, we used mutant mice expressing the active diphtheria toxin receptor subunit (DTA) exclusively in *Xcr1*-expressing cells; these mice were generated by crossing the *Xcr1C* knock-in mice (*27, 28*) with mice expressing DTA from the ubiquitous *Rosa26* locus under control of a lox-STOP-lox cassette (*R26^lsl-DTA^*) (*29*). YUMM1.7 tumors were transplanted into cohorts of male and female *Xcr1^iCre^;R26^lsl-DTA^* mice (*Xcr1^DTA^*) or DTA-negative littermate controls (*R26^lsl-DTA^* or *Xcr1^iCre^* mice; WT). Flow cytometry analysis confirmed
 that tumor-associated cDC1s were completely absent in *Xcr1^DTA^* mice, compared to WT controls (Fig.1D). The kinetics of YUMM1.7 tumor growth was unaffected in both WT and *Xcr1^DTA^* male mice. However, tumor growth was significantly increased in female *Xcr1^DTA^* mice compared to WT controls (Fig.1E,F). Thus, Xcr1^+^ cDC1s were required for control of immunogenic tumors.

Increased tumor growth upon depletion of cDC1s in female mice was associated with a drastic decrease in the recruitment of tumor-infiltrating CD8^+^ T cells (Fig.1G and Fig.S1E). Depletion of cDC1s also significantly reduced the frequency of tumor (HY)-specific CD8^+^ T cells in YUMM1.7 tumors from female mice (Fig.1H and Fig.S1F), and importantly the accumulation of IFN-γ and granzyme B (GrB)-expressing cytotoxic CD8^+^ T cells (CTLs) (Fig.1I). Interestingly, despite the significant decrease in recruitment of tumor-specific effector CD8^+^ T cells upon depletion of Xcr1^+^ cDC1s, the frequency of HY-specific CD8^+^ T cells and CTLs in tumor-draining lymph nodes (TDLN) was unaffected (Fig.S1G-I), suggesting that cDC1-mediated activation of tumor-infiltrating CD8^+^ T cells in immunogenic tumors occurs locally at the tumor site, rather than in TDLNs by migratory DCs.

To rule out potential effects of sexual dimorphism on differences in YUMM1.7 tumor growth, we used an immunogenic YUMM1.7 tumor line derived from UV-irradiated male tumor-bearing mice (YUMMER1.7) (*22*). These tumors exhibit a high somatic mutational burden and, unlike YUMM1.7, their growth was significantly decreased in immunocompetent male mice as compared to immunodeficient mice (*22*), presumably due to increased expression of neoantigens. As expected, YUMMER1.7 tumors spontaneously regressed in syngeneic male WT mice (Fig.1J,K), which was associated with the accumulation of IFN-γ and TNF-α-expressing CD8^+^ T cells in TDLNs (Fig.1L). However, male *Xcr1^DTA^* mice completely failed to control the growth of YUMMER1.7 tumors (Fig.1M,N), which was accompanied by a reduced accumulation of activated CD8^+^ T cells in TDLNs (Fig.1O,P), but not non tumor draining lymph nodes (Fig.S1J).

These experiments demonstrated the specific contribution of cDC1s to the recruitment and activation of effector T cells and the control of immunogenic tumor growth.

### Defining the molecular determinants of cDC1 maturation in tumors

The molecular pathways that control the activation and functions of tumor-infiltrating cDC1s remain largely unknown. In order to reveal the molecular determinants of intra-tumoral cDC1 maturation during control of immunogenic tumor growth, we performed single-cell RNA sequencing (scRNA-seq) of cDC1s isolated from YUMM1.7 tumors in female mice. Considering the scarcity of cDC1s in tumors, we used an index cell sorting approach which enabled us to track the cell surface phenotype of single cells and their transriptomes. We used a novel flow cytometry-based 5’-end scRNA-seq method termed FB5P-seq (*30*), which enables the sequencing of phenotypically defined single cells and incorporates unique molecular identifiers (UMIs) for accurate molecular counting. After quality control and contaminant filtering, we obtained single-cell transcriptomes from 113 *bona fide* cDC1s. Unsupervised clustering analysis using Seurat (*28*) revealed significant heterogeneity among intra-tumoral cDC1s, with 4 clusters (C0-3) visualized by Uniform Manifold Approximation and Projection (UMAP) and expressing highly distinct gene signatures (Fig.2A and Fig.S2A). Gene set enrichment analysis (GSEA) across the 4 clusters, combining gene sets from the MsigDB public repository and previsouly published DC gene signatures, revealed enrichment for distinct biological functions and activation states (Fig.2B). C3 was enriched for signatures associated with cell proliferation (E2F targets, G2M checkpoint and DNA replication) and an immature state (Core DN vu manh (*31*), Mat OFF Ardouin (*32*)) (Fig.2B,D). C0 also showed enrichment of signatures from immature cDC1s. Conversely, C1 showed a high enrichment for genes upregulated in mature cDC1s (Core UP vu manh (*31*), Mat ON Ardouin (*32*)) (Fig.2B). In addition, genes asscociated with NF-κB and IFN signaling pathways were highly enriched in C1. EnrichR analyses using KEGG, Reactome and PPI databases also revealed an enrichment for the NF-κB pathway in C1 (Fig.S2B). Accordingly, the typical DC maturation markers *Ccr7, Cd40, Fscn1,* and *Cd86,* as well as the proinflammatory cytokines *Il12b, Ccl5* and *Ccl22,* which are known target genes for the NF-κB pathway in DC(*33*), were highly expressed in C1 (Fig.2C). These data suggested NF-κB signaling to be a key regulator of intra-tumoral cDC1 maturation, which prompted us to further explore the specific role of NF-κB in cDC1s.

**Figure 2.**
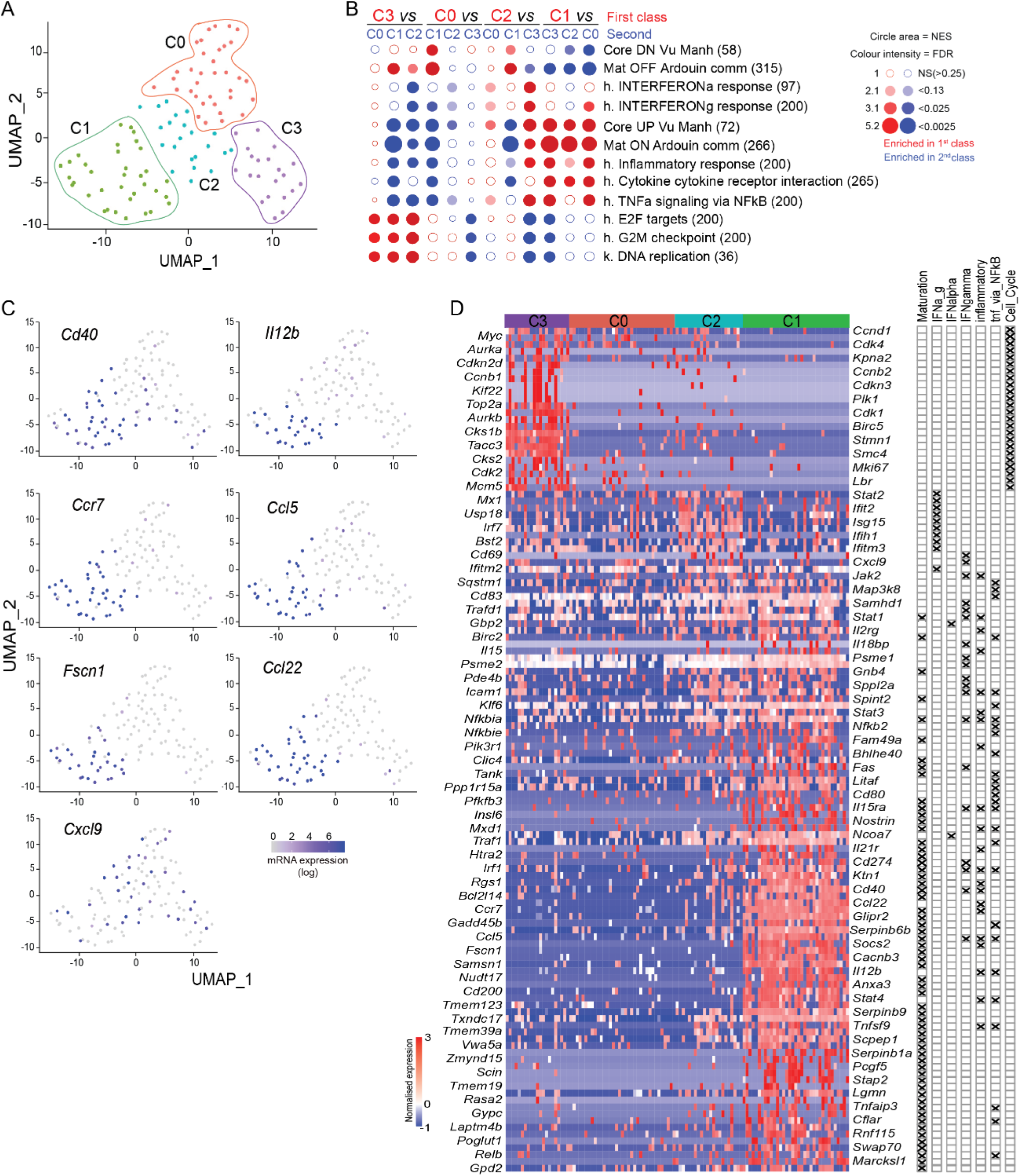
Single-cell transcriptional profiling of cDC1s from immunogenic tumors. (**A**) Dimensionality reduction performed using the Uniform Manifold Approximation and Projection (UMAP) algorithm and graph-based cell clustering using Seurat, for 113 bona fide cDC1s sorted from YUMM1.7 tumors in female mice; 4 distinct clusters are indicated C0-3. (**B**) GSEA using hallmark (h), Kegg pathway (k) and published DC maturation genesets and pairwise comparisons between clusters C0-3 in **A**, performed using BubbleMap (Bubble GUM). Circle area indicates enrichment score (NES) and color intensity false discovery rate (FDR). FDR was further corrected for multiple testing, leading to a higher stringency of the significance threshold. Core_DN_Vu_Manh and Core_UP_Vu_Manh correspond to genesets down or up-regulated, respectively, across human and mouse DC subsets during maturation in response to microbial stimuli (*31*). Mat_ON_Ardouin_comm and Mat_OFF_Ardouin_comm correspond to genesets up or down-regulated, respectively, during steady-state maturation of mouse cDC1s (*32*). Numbers in parentheses correspond to the number of genes. (**C**) Expression of selected genes related to DC maturation on the UMAP space. (**D**) Heatmap showing expression of selected genes across C0-C3. Correlations of individual genes with GSEA analysis in **B** is shown on the right.

### NF-κB controls IFN-γ-mediated programming of cDC1s in tumors

To explore the role of NF-κB activation in cDC1s, we generated mice with a conditional deletion of IKKβ (*Ikbkb*), a critical kinase for NF-κB activation, specifically in cDC1s. We crossed *Xcr1^iCre^* mice with *Ikbkb*^F/F^ mice and confirmed efficient deletion of IKKβ specifically in cDC1s in homozygous *Ikbkb*^F/F^ *Xcr1^iCre/+^* progeny (*Ikbkb^ΔXcr1^),* at both mRNA and protein levels (Fig.S3A-E). We next transplanted YUMM1.7 tumors in cohorts of male and female *Ikbkb*^ΔXcr1^ mice or littermate *Ikbkb*^F/F^ controls. IKKβ deletion in cDC1s had no impact on tumor growth in male mice (Fig.S3F,G). However, YUMM1.7 tumor growth was significantly increased in *Ikbkb^ΔXcr1^* female mice compared with littermate controls (Fig.3A). In addition, *Ikbkb^ΔXcr1^* male mice failed to control the growth of YUMMER1.7 tumors (Fig.3B). These results demonstrated that IKKβ was required in cDC1s for control of immunogenic tumor growth. IKKβ deletion did not affect cDC1 accumulation in tumors (Fig.S3H), but reduced their expression of the co-stimulatory molecules CD40, CD86 and CCR7 (Fig.3C,D), showing that cell-intrinsic NF-κB signaling was required for optimal maturation of tumor-associated cDC1s. Similarly to *Xcr1^DTA^* mice, female *Ikbkb^ΔXcr1^* mice showed a strong decrease in the recruitment and activation of CD8^+^ T cells in YUMM1.7 tumors (Fig.3E,F and Fig.S3I-K). Furthermore, the frequency of HY-specific CD8^+^ T cells was significantly reduced in tumors from *Ikbkb*^ΔXcr1^ mice (Fig.3G and Fig.S3L), in line with data from *Xcr1^DTA^* mice. The impaired activation state of tumor-infiltrating CD8^+^ T cells in *Ikbkb^ΔXcr1^* mice was confirmed by gene expression analysis showing reduced expression of several T cell activation markers^21,22^ (Fig.3H). In the case of YUMMER1.7 tumors, increased tumor growth was associated with reduced expression of maturation markers by cDC1s and decreased accumulation of activated CD8^+^ T cells in TDLNs (Fig.3I,J), but not in non-draining LNs (Fig.S3M).

**Figure 3.**
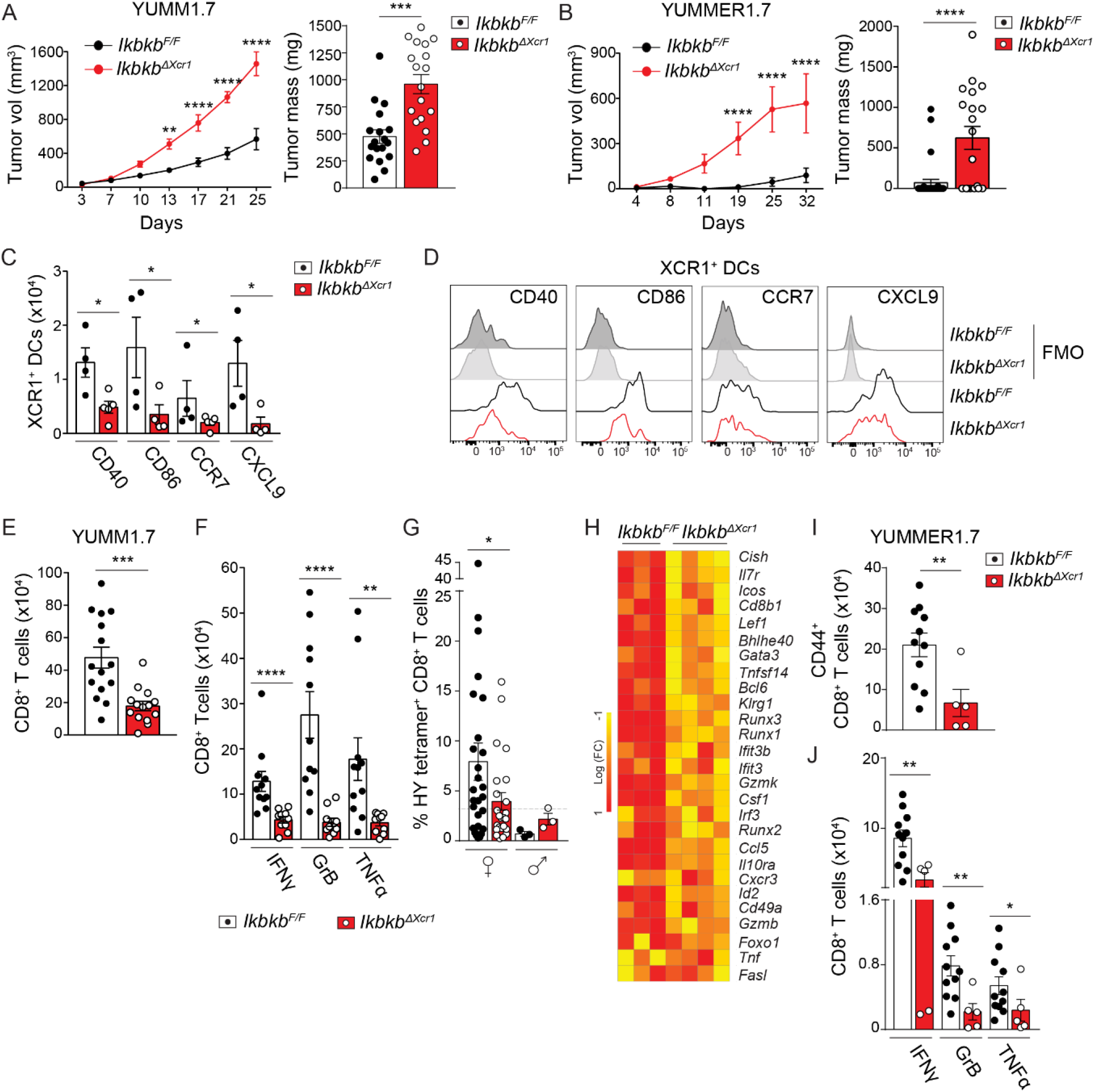
IKKβ in cDC1s is required for control of immunogenic tumors. (**A**) YUMM1.7 tumor growth in female *Ikbkb^F/F^* (n=11) and *Ikbktf^Xcr1^* (n=9) mice. (**B**) YUMMER1.7 tumor growth in male *Ikbkb^F/F^* (n=33) and *Ikbkb^ΔXcr1^ f*n=19) mice. (**C**) Flow cytometry analysis of tumor-infiltrating cDC1s from YUMM1.7 tumors in *Ikbkb^F/F^* and *Ikbkb^ΔXcr1^* female mice; total numbers of CD40, CD86, CCR7 and CXCL9-expressing cDC1s are shown. (**D**) Representative histograms of flow cytometry analysis in **C**, including FMO control. (**E**) Total number of CD8^+^ T cells and (**F**) numbers of IFN-γ, GrB or TNF-α expressing cells in YUMM1.7 tumors from female *Ikbkb^F/F^* and *Ikbkb^ΔXcr1^* mice. (**G**) Frequency of HY-specific tumor-infiltrating CD8^+^ T cells in YUMM1.7 tumors from male and female *Ikbkb^F/F^* and *Ikbkb^ΔXcr1^* mice. (**H**) Differential expression of T cell activation genes by CD8^+^ T cells from YUMM1.7 tumors in female *Ikbkb^F/F^* and *Ikbkb^ΔXcr1^* mice. (**I**) Total numbers of CD44^+^ CD8^+^ T cells in TDLN from *Ikbkb^F/F^* and *Ikbkb^ΔXcr1^* male mice bearing YUMMER1.7 tumors. (**J**) Numbers of IFN-γ, GrB and TNF-α expressing CD44^+^ CD8^+^ T cells in TDLN from male YUMMER1.7 tumor-bearing mice. Total cell numbers are indicated for 250mg of tumor tissue. Pooled or representative data are shown from at least 2 independent experiments. Data are shown as mean ± SEM. In scatter plots each point corresponds to an individual mouse. Statistical analysis was performed by two-way ANOVA followed by Sidak’s multiple comparison tests (**A** and **B**) or Mann-Whitney test. Fisher’s exact test was used for frequency of tetramer positive cells when their proportion > the highest level obtained with HPV16 E7_(49-57)_/H-2Db tetramer (**G**).

Production of the chemokine CXCL9 by DCs has been suggested to play a key role in the recruitment of effector CD8^+^ T cells to tumors (*1*). We observed a significant reduction in CXCL9-expressing cDC1s in *Ikbkb^ΔXcr1^* mice compared to littermate controls (Fig.3C,D). In addition, uncontrolled growth of YUMMER1.7 tumors in *Ikbkb*^ΔXcr1^ mice was also associated with impaired expression of CXCL9 by cDC1s in TDLNs (Fig.S3N,O), which correlated with reduced accumulation of activated CD8^+^ T cells (Fig.3I,J).

These data demonstrated that NF-κB activation in intra-tumoral cDC1s was required for the recruitment and activation of CD8^+^ T cells and control of immunogenic tumors, likely via the promotion of cDC1 functional maturation, including their expression of CXCL9.

To gain further insight into the molecular mechanisms by which NF-κB drives cDC1-mediated control of tumor growth, we performed bulk RNA-seq analysis of cDC1s sorted from YUMM1.7 tumors in female *Ikbkb*^F/F^ and *Ikbkb^ΔXcr1^* mice. GSEA revealed that IFN pathway gene signatures were highly enriched in wild-type cDC1s (*Ikbkb*^F/F^) compared to IKKβ-deficient cells (*Ikbkb^ΔXcr1^*) (Fig.4A), with a particularly strong negative enrichment for the *IFN_gamma_response* pathway (Fig.4B). As expected, the *TNFA_signaling_via_NF-κB* pathway was also strongly enriched in wild-type cells, confirming an impaired activation of NF-κB after IKKβ deletion (Fig.4A). To confirm changes in the IFN-γ response program in intra-tumoral cDC1s, we measured by qRT-PCR the expression levels of hallmark genes induced by IFN-γ(*34*). IKKβ deletion in cDC1s led to a consistent down-regulation of most of these genes, confirming the NF-κB dependent control of the IFN-γ response in intra-tumoral cDC1s (Fig.4C). This prompted us to assess the impact of IFN-γ signaling on cDC1 functionality during immunogenic tumor growth. For this purpose, we treated female YUMM1.7 tumor-bearing mice with IFN-γ-neutralizing antibody. This resulted in a significant increase in tumor growth (Fig.4D,E) and reduction in tumor-infiltrating effector CD8^+^ T cells (Fig.4F). We then measured the expression of hallmark genes induced by IFN-γ in cDC1s isolated from these tumors. Similarly to conditional deletion of IKKβ in cDC1s, IFN-γ neutralization reduced the expression of most of these genes (Fig.4G), which confirmed the association between NF-κB activation and induction of IFN-γ-responsive genes in intra-tumoral cDC1s.

**Figure 4.**
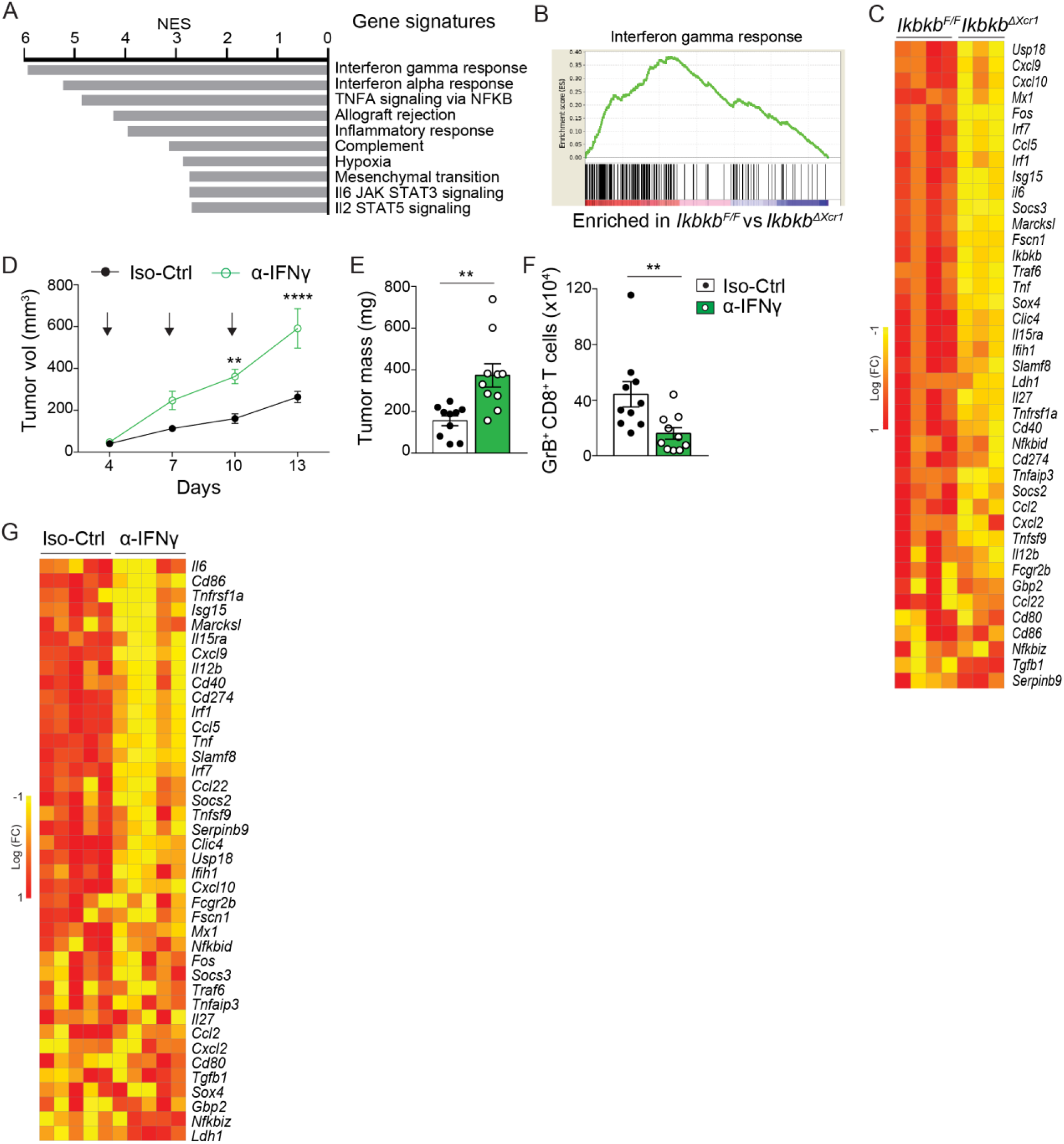
IKKβ controls IFN-γ-mediated programming of intra-tumoral cDC1s. (**A**) GSEA of bulk RNA-seq data from cDC1s isolated from YUMM1.7 tumors in female *Ikbkb^F/F^* and *Ikbkb^ΔXcr1^* mice. 10 genesets with highest enrichment in cDC1s from wild-type (*Ikbkb*^F/F^) mice as compared to *Ikbkb^ΔXcr1^* mice are shown. (**B**) Enrichment plot for Interferon_gamma_response in cDC1s from YUMM1.7 tumors in *Ikbkb^F/F^* and *Ikbkb^ΔXcr1^* female mice. (**C**) Differential expression of IFN-regulated genes in cDC1s from YUMM1.7 tumors in female *Ikbkb^F/F^* and *Ikbkb^ΔXcr1^* mice. (**D**,**E**) YUMM1.7 tumor growth in female mice treated with anti-IFN-γ (α-IFNγ) or isotype control (Iso-Ctrl) at indicated time points (arrows). In (**D**), n=10 for α-IFNγ and Iso-Ctrl treated mice. (**F**) Total numbers of GrB-expressing CD8^+^ T cells in YUMM1.7 tumors from female mice after treatment with anti-IFN-γ (per 250mg tumor tissue). (**G**) Differential expression of selected IFN-regulated genes in cDC1s from YUMM1.7 tumors in female mice after treatment with anti-IFN-γ. Representative data are shown from at least 2 independent experiments. Data are shown as mean ± SEM. In scatter plots each point corresponds to an individual mouse. Statistical analysis was performed with two-way ANOVA followed by Sidak’s multiple comparison tests (**D**) or Mann-Whitney test (**E** and **F**).

NF-κB/IFN-γ-dependent genes in tumor-associated cDC1s included both *Cxcl9* and *Cxcl10* (Fig.4C,G), which are major chemoattractants for CXCR3-expressing effector CD8^+^ T cells (*1, 36, 37*). To evaluate the contribution of these CXCR3-ligands to the control of tumor growth, we treated YUMM1.7 tumor-bearing female mice with CXCR3-blocking antibody. As observed with IKKβ deletion in cDC1s and with IFN-γ neutralization, CXCR3-blockade impaired the recruitment of tumorinfiltrating CD8^+^ T cells and increased tumor growth (Fig.S4). These results support the hypothesis that control of tumor growth by cDC1s was indeed dependent on their production of chemokines promoting the intra-tumoral recruitment of CD8^+^ T cells in a CXCR3-dependant manner.

### NF-κB mediated IRF1 expression in cDC1s is required for control of immunogenic tumors

Among the NF-κB and IFN-γ regulated genes in tumor-associated cDC1s was the transcription factor IRF1 (Fig.4C,G), which is a master regulator of IFN-mediated gene expression (*38*) and also enriched during intra-tumoral cDC1 maturation (Fig.S2C). We next sought to determine the role of IRF1 in the NF-κB dependent control of cDC1-mediated anti-tumor immunity. To this end, we generated mice with a conditional deletion of IRF1 in cDC1s (*Irf1^ΔXcr1^*). We confirmed efficient deletion of IRF1 expression specifically in cDC1s from *Irf1^ΔXcr1^* mice at mRNA and protein levels (Fig.S5A-D). We then transplanted cohorts of male and female *Irf1^ΔXcr1^* mice or littermate controls with YUMM1.7 tumors, or only male mice with YUMMER1.7 tumors. As we had observed in *Xcr1^DTA^* and *Ikbkb^ΔXcr1^* mice, growth of YUMM1.7 tumors was unaffected in male *Irf1^ΔXcr1^* mice (Fig.S5E), but was significantly increased in female *Irf1^ΔXcr1^* mice compared to controls (Fig.5A). Similarly, *Irf1^ΔXcr1^* male mice completely failed to control growth of YUMMER1.7 tumors (Fig.5B). In line with data from *Ikbkb^ΔXcr1^* mice, the accumulation of cDC1s in YUMM1.7 tumors from female *Irf1^ΔXcr1^* mice was unaffected (Fig.S5F), but their expression of maturation markers and CXCL9 was impaired (Fig.5C). Furthermore, the expression of IFN-γ responsive genes was down-regulated in intra-tumoral cDC1s from *Irf1^ΔXcr1^* mice (Fig.5D). Consistent with the increased growth of YUMM1.7 tumors in female *Irf1^△χc^?~?* mice, the frequency of tumor-infiltrating HY-specific CD8^+^ T cells was significantly reduced (Fig.5G and Fig.S5G), as was the global recruitment and activation of tumor-infiltrating effector CD8^+^ T cells (Fig.5E-H and Fig.S5H). Similarly, the frequency of activated CD8^+^ T cells in TDLNs after engraftment with YUMMER1.7 tumors was significantly reduced in *Irf1^ΔXcr1^* mice (Fig.5I,J).

**Figure 5.**
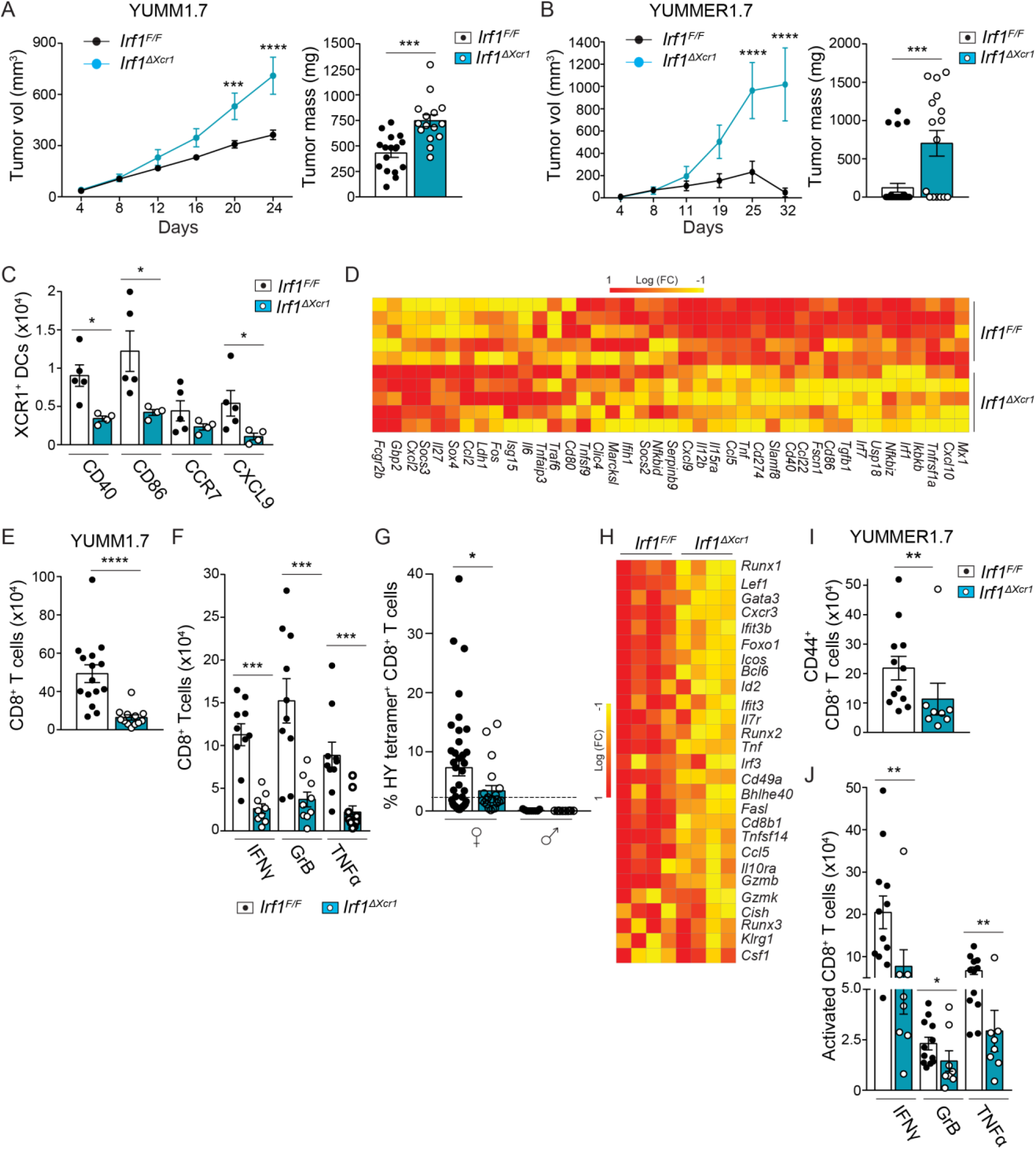
IRF1 in cDC1s is required for control of immunogenic tumors. (**A**) YUMM1.7 tumor growth in female *Irf1^F/F^* (n=14) and *Irf1^ΔXcr1^* (n=9) mice. (**B**) YUMMER1.7 tumor growth in male *Irf1^F/F^* (n=34) and *Irf1^ΔXcr1^* (n=16) mice. (**C**) Flow cytometry analysis of tumor-infiltrating cDC1s from YUMM1.7 tumors in *Irf1^F/F^* and *Irf1^ΔXcr1^* female mice; total numbers of CD40, CD86, CCR7 and CXCL9-expressing cDC1s are shown. (**D**) Differential expression of IFN-regulated genes in cDC1s from YUMM1.7 tumors in female *Irf1^F/F^* and *Irf1^ΔXcr1^* mice. (**E**) Total number of CD8^+^ T cells and (**F**) numbers of IFN-γ, GrB or TNF-α expressing cells in YUMM1.7 tumors from female *Irf1^F/F^* and *Irf1^ΔXcr1^* mice. (**G**) Frequency of HY-specific tumor-infiltrating CD8^+^ T cells in YUMM1.7 tumors from male and female *Irf1^F/F^* and *Irf1^ΔXcr1^* mice. (**H**) Differential expression of T cell activation genes by CD8^+^ T cells from YUMM1.7 tumors in female *Irf1^F/F^* and *Irf1^ΔXcr1^* mice. (**I**) Total numbers of CD44^+^ CD8^+^ T cells in TDLN from *Irf1^F/F^* and *Irf1^ΔXcr1^* male mice bearing YUMMER1.7 tumors. (**J**) Numbers of IFN-γ, GrB and TNF-α expressing CD44^+^ CD8^+^ T cells in TDLN from male YUMMER1.7 tumorbearing mice. Total cell numbers are indicated for 250mg of tumor tissue. Pooled or representative data are shown from at least 2 independent experiments. Data are shown as mean ± SEM. In scatter plots each point corresponds to an individual mouse. Statistical analysis was performed by two-way ANOVA followed by Sidak’s multiple comparison tests (**A** and **B**) or Mann-Whitney test. Fisher’s exact test was used for frequency of tetramer positive cells when their proportion > the highest level obtained with HPV16 E7(49-57)/H-2Db tetramer (**G**).

These data demonstrated that IRF1, along with NF-κB, was required for the cDC1-mediated control of immunogenic tumors. Furthermore, IRF1 was necessary for the induction of IFN-γ-responsive genes in intra-tumoral cDC1s down-stream of NF-κB activation, identifying a hierarchical relationship between NF-κB and IRF1 activation in cDC1s which is required for control of immunogenic tumor growth.

### NF-κB and IRF1 coordinate maturation of tumor-infiltrating cDC1s

To further dissect the impact of NF-κB and IRF1-regulated pathways on the intra-tumoral maturation of cDC1s, we performed scRNA-seq analysis on cDC1s from YUMM1.7 tumor-bearing mice after targeted deletion of IKKβ or IRF1. We analysed a total of 361 single cell transcriptomes of index-sorted *bona fide* cDC1s, using FB5P-seq, and performed unsupervised Seurat clustering analysis. We identified 7 distinct cDC1 clusters based on their gene expression (Fig.6A,B). GSEA revealed enrichment of cell proliferation gene signatures in C5, and to a lesser extent C6 (Fig.6C,D), which was similar to C3 in the previous analysis of wild-type cDC1s (WT-C3) (Fig.2A,B and Fig.S6B). C1 and C2 were enriched for inflammatory pathways and DC maturation gene modules (Fig.6C,D and Fig.S6A), representing the counterparts of the mature cDC1s identified among wild-type cDC1s (WT-C1) (Fig.S6B). In contrast, C3 was enriched for immature DC genesets and expressed the lowest levels of DC maturation genes (Fig.6C,D and Fig.S6A), mirroring the cluster of immature WT cDC1s identified previously (WT-C0 in Fig.2) (Fig.S6B).

**Figure 6.**
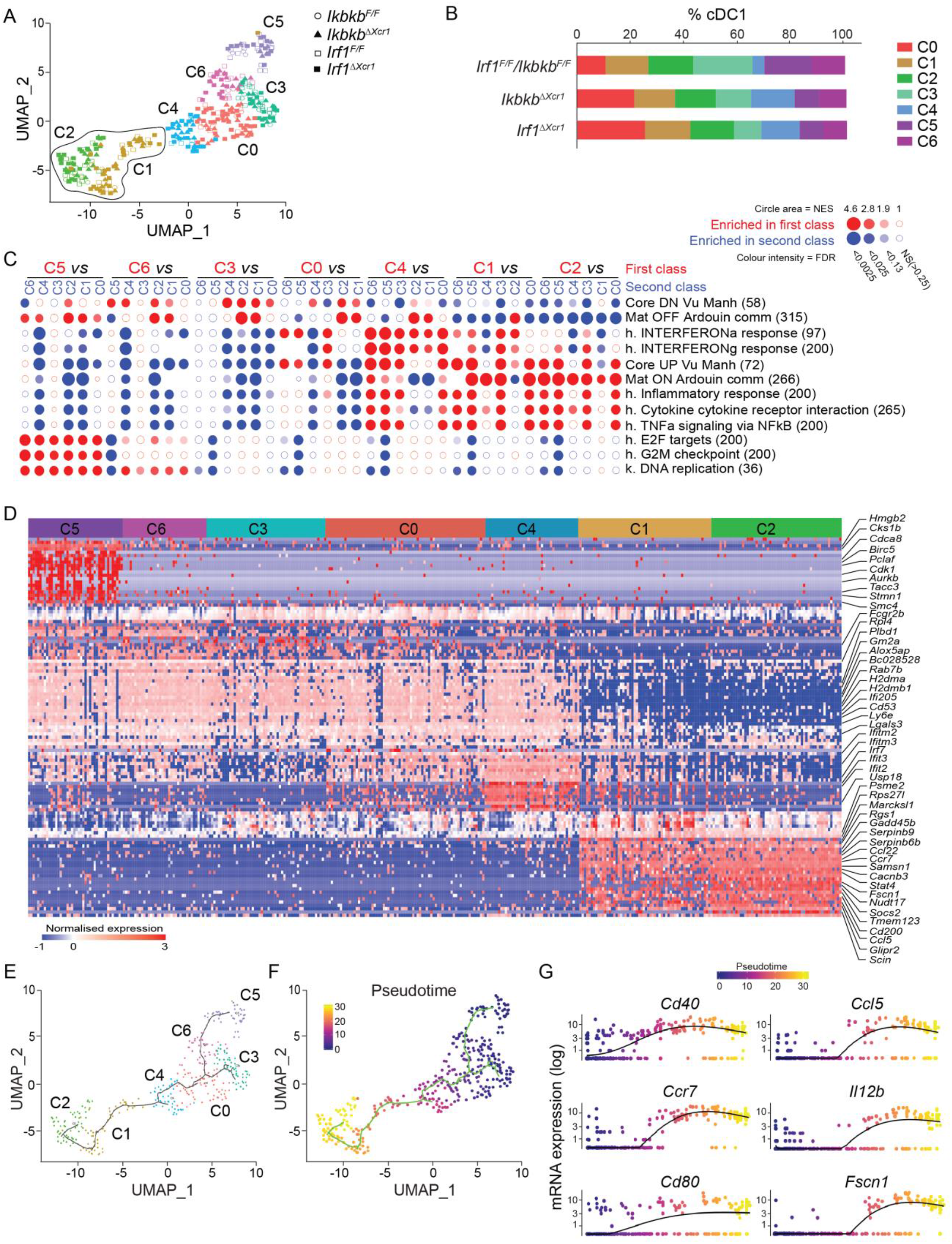
IKKβ and IRF1 coordinate intra-tumoral cDC1 maturation. (**A**) UMAP and graph-based cell clustering for 360 bona fide cDC1s isolated from YUMM1.7 tumors in female *Ikbkb^F/F^, Ikbkb^ΔXCR1^, Irf1^F/F^* and *Irf1^ΔXcr1^* mice; 7 distinct clusters are indicated C0-6. (**B**) Distribution of clusters among total cDC1s in each group. (**C**) GSEA using hallmark (h), Kegg pathway (k) and published DC maturation genesets and pairwise comparisons between clusters C0-6 in **A**, performed using BubbleMap (Bubble GUM) as described in figure 2. (**D**) Heatmap showing expression of selected genes across C0-C6. (**E,F**) Single-cell trajectory inference by Monocle and projection onto the UMAP space; (**E**) showing Seurat clusters and (**F**) the pseudotime analysis using C5 as the root. (**G**) Gene expression profile of selected genes related to cDC1 maturation in pseudotime calculated by Monocle.

To evaluate the possible dynamics between these different clusters, we performed single-cell trajectory inference using Monocle (*39*). This unsupervised learning algorithm inferred a branched trajectory of cDC1s with three end points at C5, C3 and C2 (Fig.6E). Since C2 expressed the highest enrichment for DC maturation genes (Fig.6C), which were negatively enriched in C3 and C5, we considered C3 and C5 as alternative roots for a cDC1 maturation endpoint at C2 (Fig.6E). These two alternative paths for cDC1 maturation converged at C4, before reaching the fully mature state represented by C2 (Fig.6E-G). This analysis pinpointed C4 as a possible transitory state for intra-tumoral cDC1 maturation. C4 showed a particular enrichment for IFN-responsive genes, as well as an induction of DC maturation genesets (Fig.6C). Interestingly, the proportion of cells in C4 substantially increased upon deletion of NF-κB or IRF1 (Fig.6B), pointing to a possible role for NF-κB/IRF1 signaling in promoting the exit of cDC1s from this IFN-responsive state (C4) into their fully mature state (C2). In keeping with this hypothesis, both NF-κB signaling and IRF1 expression were mostly enriched in fully mature cDC1s (C1 and C2) (Fig.6C and Fig.S6C). Furthermore, the genes upregulated in C1 and C2 compared to C4, including *Ccl5, Il12b, Cd40* and *Fscn1* (Fig.S6A,D), were down-regulated in both NF-κB and IRF1 deficient intra-tumoral cDC1s (Fig.4C, Fig.5D and Fig.S6D). The transcripts down-regulated in intra-tumoral cDC1s upon IKKβ and IRF1 deletion included genes associated with antigen-processing and presentation (e.g., *Tapbp* and *Psmb1),* as well as DC maturation and IFN response pathways (Fig.S6D,E). These data support a role for NF-κB/IRF1 signaling in promoting the functional maturation of intra-tumoral cDC1s, including antigen-presentation as well as cytokine and chemokine production leading to recruitment and activation of effector T cells.

Moreover, consistent with IRF1 acting down-stream of NF-κB activation, EnrichR analysis of DEGs identified upon IKKβ deletion in cDC1s using transcription factor PPI database revealed a network including three members of the IRF family that emerged with high combined scores, of which IRF1 had the highest number of interactions within the network (Fig.S6H,I).

Taken together, these data show that an NF-κB/IRF1 axis coordinates the maturation of tumorinfiltrating cDC1s, and we have pinpointed a transitional activation state which is highly dependent on NF-κB and IRF1 signaling for progression to a fully mature functional state.

In order to extend our observations to other tumor models, we analysed the expression of NF-κB/IRF1 co-dependent genes in scRNA-seq data from cDC1s in a previous study from a mouse model of lung adenocarcinoma (*15*). We first reconstructed Seurat clustering of cDC1s from naïve and tumor-bearing lungs in this dataset (gene expression omnibus (GEO) accession GSE131957). This identified 8 distinct clusters visualized by UMAP (C0-7) (Fig.S7A). We next performed a Jaccard similarity analysis to compare these clusters with the cDC1 clusters we identified in YUMM1.7 melanomas (YUMM-C0/C6). The highest Jaccard index was obtained between C7 of lung tumor-associated cDC1s and YUMM-C1/C2 mature cDC1s (Fig.S7B), suggesting a similar maturation state between tumor-associated cDC1s in melanomas and lung adenocarcinomas. Among lung cDC1s, IKKβ and IRF1 were also most highly expressed by cells with a mature phenotype (C7), as compared to any other cluster (Fig.S7D). This was accompanied by the highest expression levels of NF-κB/IRF1*-*dependent genes involved in IFN-γ response and DC maturation. Of note, mature cDC1s in normal mouse lung tissue did not show a concerted up-regulation of IKKβ and IRF1 (Fig.S7E), suggesting that NF-κB and IRF1 activation in cDC1s is up-regulated in response to specific signals in the tumor-microenvironment.

In summary, we have described in a clinically relevant mouse model of melanoma the coordinated regulation of intra-tumoral cDC1 maturation by NF-κB and IRF1, that is required to control immunogenic tumor growth. In addition, the implication of the NF-κB/IRF1 axis in intra-tumoral cDC1 maturation appears to be conserved in other cancer models.

### The NF-κB/IRF1 axis in cDC1s correlates with good prognosis in cancer patients

We next sought to determine the relevance of NF-κB and IRF1 signaling in cDC1s to human cancer. We first deconvoluted the gene expression data from The Cancer Genome Atlas (TCGA) for skin cutaneous melanoma (SKCM, 468 patients). We used a previously published gene list for cDC1s from this dataset (*40*) and scored the expression of these genes in addition to *IRF1* and two hallmark transcripts of the NF-κB pathway (*IKBKB* and *NFKB1*) (Fig.7A). Hierarchical clustering showed consistency between *IRF1, IKBKB* and *NFKB1* and the human cDC1 gene signature, therefore we included these genes in the cDC1 signature for inference of NF-κB/IRF1 pathway enrichment in cDC1s. To derive the most appropriate transcripts for the identification of activated CD8^+^ T cells, we selected the most relevant genes specific to CD8^+^ T cells that were altered upon IKKβ or IRF1 deletion in cDC1s in our mouse melanoma model (Fig.7B). In line with the established role of cDC1s in recruitment of tumor-infiltrating CD8^+^ T cells, the cDC1 signature showed a high degree of positive correlation with the activated CD8^+^ T cell signature (Fig.7C). To discern whether this observation was relevant for disease prognosis, we analysed the survival of melanoma patients with high (top quartile) and low (bottom quartile) levels of cDC1 alone and NF-κB/IRF1-enriched cDC1 signatures. We found that the NF-κB/IRF1-enriched cDC1 signature was associated with improved prognostic outcome in melanoma patients compared to the cDC1 signature alone (Fig.7D).

**Figure 7.**
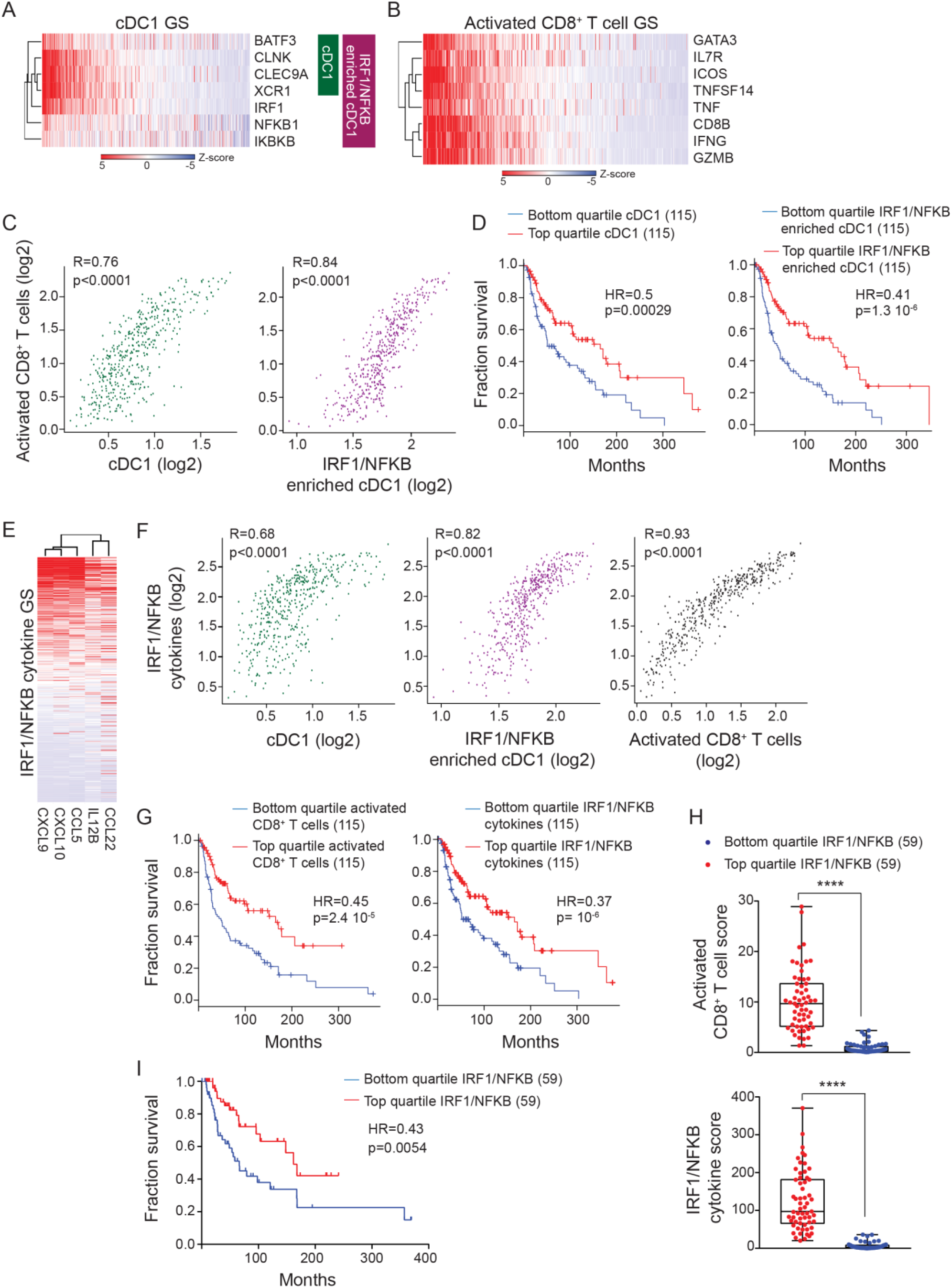
NF-κB/IRF1 enriched cDC1 gene signatures correlate with increased survival in melanoma patients. (**A**) Heatmap showing the gene-similarity-ordered, z-transformed expression values for cDC1 and NFKB/IRF1-enriched cDC1 gene signatures (GS) and (**B**) activated CD8^+^ T cell GS in melanoma patients from TCGA datasets. (**C**) Spearman correlation between cDC1 GS or NFKB/IRF1-enriched cDC1 GS with activated CD8^+^ T cell GS in melanoma patients from TCGA datasets. (**D**) Kaplan-Meier overall survival curves of TCGA melanoma patients comparing top and bottom quartile of cDC1 GS or NFKB/IRF1-enriched cDC1 GS. Hazard ratio (HR) and *p* value from univariate cox regression are indicated. (**E**) Heatmap showing the gene-similarity-ordered, z-transformed expression values for NFKB/IRF1-dependent chemokine GS in melanoma patients from TCGA datasets. (**F**) Spearman correlation between cDC1 GS or NFKB/IRF1-enriched cDC1 GS or activated CD8^+^ T cell GS with NFKB/IRF1-dependent chemokine GS in melanoma patients from TCGA datasets. (**G**) Kaplan-Meier overall survival curves of TCGA melanoma patients comparing top and bottom quartile of activated CD8^+^ T cell GS or NFKB/IRF1-dependent chemokine GS. (**H**) Box plots comparing activated CD8^+^ T cell GS and NFKB/IRF1-dependent chemokine GS in top and bottom quartile of NFKB/IRF1 expression in TCGA melanoma patients bearing high cDC1 GS (top 50%). (**I**) Kaplan-Meier overall survival curves comparing top and bottom quartile of NFKB/IRF1 expression in TCGA melanoma patients bearing high cDC1 GS score (top 50%).

To further explore the implication of the NF-κB/IRF1 axis in cDC1s for human melanoma, we analysed the expression of NF-κB/IRF1 co-regulated cytokine genes that we had identified in cDC1s from our mouse melanoma model, and which were also linked to CD8^+^ T cell recruitment and activation (*CCL5, CCL22, CXCL9, CXCL10* and *IL12B*) (Fig.7E). These cytokines are highly expressed in activated cDCs compared to other immune cells in human cancers^40^. This NF-κB/IRF1-dependent cytokine signature was highly correlated with the activated CD8^+^ T cell signature and showed a higher correlation with the NF-κB/IRF1-enriched cDC1 signature than with the cDC1 signature alone (Fig.7F), supporting a role for the NF-κB/IRF1 axis in expression of these cytokines and recruitment of cytotoxic CD8^+^ T cells in melanoma patients. Consistent with this observation, high expression of both the activated CD8^+^ T cell signature and the NF-κB/IRF1-dependent cytokine signature, were significantly associated with patient survival (Fig.7G). To account for cDC1-independent NF-κB/IRF1 expression in our analysis, that is more likely in melanoma patients bearing a low cDC1 score, we focused our analysis on data from patients bearing high cDC1 gene signatures. In this analysis, we compared the score of activated CD8^+^ T cells and NF-κB/IRF1-dependent cytokines between individuals with high (top quartile) and low (bottom quartile) levels of *IRF1* and *IKBKB/NFKB1* gene expression. This revealed that among patients bearing high cDC1 signatures, individuals with higher expression of *IKBKB*/*NFKB1* and *IRF1* showed a substantial increase in the CD8^+^ T cell signature and NF-κB/IRF1-dependent cytokine signature (Fig.7H). Furthermore, *IKBKB*/*NFKB1* and *IRF1* expression correlated significantly with good prognosis for melanoma patients bearing high cDC1 signatures (Fig.7I). Although we cannot ascertain from this analysis that NF-κB/IRF1-dependent genes are exclusively or even dominantly expressed by intra-tumoral cDC1s, these findings point to a powerful influence of NF-κB and IRF1 in melanoma patients and the interplay between tumor-associated cDC1s and CD8^+^ T cells that are critical factors in prognosis.

## Discussion

Several studies in preclinical mouse models and correlative analyses in cancer patients support an important contribution of cDC1s to the efficacy of immunotherapy. However, the roles of cDC1s in relation to intrinsic tumor immunogenicity are less clear. Furthermore, the intracellular pathways that regulate cDC1 function in anti-tumor immunity are not known. Here, we examined the specific role of cDC1s in control of tumor growth in relation to tumor immunogenicity in a clinically relevant mouse melanoma model. We demonstrated that specific depletion of Xcr1-expressing cDC1s did not affect growth of non-immunogenic tumors, where activated CD8^+^ T cells were sparce, but severely impacted the control of immunogenic tumors, at least in part by preventing the recruitment of tumorspecific CD8^+^ T cells.

To further explore the phenotype of intra-tumoral Xcr1^+^ cDC1s we performed single-cell RNA sequencing. This experiment revealed significant heterogeneity among tumor-associated cDC1s and we identified distinct clusters of cells expressing genes associated with proliferation and DC maturation. Strikingly, a cluster of cDC1s in tumors expressed enrichment for genes associated with DC maturation alongside NF-κB and IFN-regulated signaling pathways, indicating that these pathways may have an important role in intra-tumoral cDC1 maturation. In previous studies, CXCL9 expression by DCs was suggested to play a key role in the recruitment of effector CD8^+^ T cells to tumors (*1*). However, CXCL9 and CXCL10 were not selectively expressed by mature cDC1s in our study. In contrast, mature cDC1s selectively up-regulated IL-12 and CCL5 expression. In another recent study, increased IL-12 production by intra-tumoral DCs was shown to be dependent on sensing of IFN-γ in mice treated with immune-checkpoint inhibitors (*16*). We showed that targeted deletion of IKKβ impaired intra-tumoral cDC1 maturation and expression of IFN-γ-regulated genes, which was associated with increased tumor growth. These data suggested an intrinsic role of NF-κB signaling in the IFN-γ-mediated licencing of cDC1s and control of tumor growth. We identified IRF1, which is a known regulator of IFN response pathways, as an NF-κB dependent gene in intra-tumoral cDC1s. Moreover, the targeted deletion of IRF1 in Xcr1^+^ cDC1s phenocopied the deletion of IKKβ, providing genetic evidence for an NF-κB/IRF1 axis regulating the maturation of intra-tumoral cDC1s and control if immunogenic tumor growth.

NF-κB and IFN-regulated pathways have been reported to converge in the regulation of a common set of genes. For example, NF-κB and ISGF3 (STAT 1, STAT2 and IRF9) cooperate downstream of IFN-I signaling in macrophages (*42, 43*). However, this has not been studied in DCs. We described a subset of intra-tumoral cDC1s distinguished by high expression of IL-12 and CCL5, as well as core DC maturation markers such as CCR7, CD40 and Fascin 1 (*Fscn1),* and expression of these genes were both NF-κB and IRF1-dependent. Since IRF1 gene expression was down-regulated upon IKKβ deletion in cDC1s, this implied that IRF1 may act down-stream of NF-κB activation. In fact, the regulatory regions of the mouse and human IRF1 genes contain 23 and 37 NF-κB binding sites, respectively (*44*). Our studies showed that NF-κB activation alone was not sufficient for full maturation of intra-tumoral cDC1s, which also required exposure to IFN-γ, associated with the recruitment of activated CD8^+^ T cells. Therefore, it is likely that IFN-γ signaling directs IRF1-mediated maturation of cDC1s. In keeping with this hypothesis, IRF1 expression was abrogated in cDC1s by neutralisation of IFN-γ, as was the expression of IRF1-dependent genes. IRF1 has previously been shown to amplify IFN-γ-mediated responses in macrophages to promote anti-microbial activity (*45*).

NF-κB dependent IRF1 expression may serve a similar function in tumor-associated cDC1s. Moreover, comparison between mouse melanoma and lung adenocarcinoma datasets showed that they shared enrichment of the same NF-κB/IRF1-responsive gene module in mature cDC1s, indicating that this pathway is conserved in other cancer models.

We used scRNA-seq data to decipher the trajectory of intra-tumoral cDC1 maturation, which revealed a transitional cell state enriched for IFN response pathways that preceded full maturation. This transitional cDC1 population increased upon NF-κB or IRF1 deletion, consistent with a role for NF-κB/IRF1 signaling in the IFN-dependent progression of cDC1 maturation. Of note, accumulation of this IFN pathway-enriched transitional state in the absence NF-κB/IRF1 signaling, implied that only a subset of IFN-responsive genes in cDC1s are required for anti-tumor immunity, and that these genes are critically regulated by NF-κB and IRF1. This is consistent with a previous study that showed a selective role of IRF1 in expression of inflammatory but not anti-viral genes during IFN responses (*38*). The enrichment of IFN response pathways in cDC1s upon deletion of NF-κB /IRF1 may also reflect cross-regulation between IFN-γ and IFN-I regulated pathways i.e, the arrest of IFN-γ-mediated cDC1 maturation upon deletion NF-κB and IRF1 may lead to a compensatory increase in cDC1 responses to IFN-I. In fact, previous studies have shown that IFN-γ and IFN-I pathways can antagonize one another in certain contexts (*46–48*).

Among the genes commonly regulated by NF-κB and IRF1 in intra-tumoral cDC1s, T cell chemokines were conspicuous, including *Ccl5, Ccl22, Cxcl9 and Cxcl10.* Interestingly, these chemokines appeared to be selectively expressed during cDC1 maturation; CXCL9 and CXCL10 were mostly expressed during the transitional cDC1 maturation state, associated with high enrichment for IFN response pathways, whereas CCL5 and CCL22 were selectively expressed in fully mature cDC1s. This was in keeping with the poor infiltration of CD8^+^ T cells in tumors upon conditional deletion of NF-κB or IRF1 signaling in cDC1s. Other NF-κB/IRF1-regulated genes in intra-tumoral cDC1s were directly related to antigen cross-presentation (*49, 50*) and T cell activation (*31, 32, 35*), suggesting that the impaired accumulation of tumor antigen-specific cytotoxic CD8^+^ T cells after deletion of NF-κB or IRF1 in cDC1s may be due in part to impaired cross-presentation of tumor antigens. Therefore, activation of the NF-κB/IRF1 axis in intra-tumoral cDC1s likely promoted both the priming of CD8^+^ T cells and their recruitment to immunogenic tumors. Additionally, cDC1 maturation in tumors was associated with NF-κB/IRF1-dependent up-regulation of IL-12 expression. In contrast to IFN response pathways, the IL-12 signaling pathway was selectively enriched among NF-κB/IRF1-dependent genes in mature cDC1s. Considering with the well-established role of IL-12 in promoting cytotoxic T cell responses, the NF-κB /IRF1-dependent expression of IL-12 by mature intra-tumoral cDC1s is likely to be critical for maintaining T cell effector functions in the tumormicroenvironment.

In summary, our data describe an NF-κB/IRF1 axis that governs intra-tumoral cDC1 maturation and their ability to control tumor growth through the recruitment and activation of CD8^+^ T cells in a clinically relevant model of melanoma. Finally, we analysed data from human melanoma in TCGA, and found a correlation between the enrichment of NF-κB/IRF1-regulated genes associated with cDC1s and activated CD8^+^ T cell genes, both of which indicated a good prognosis for patients. This suggests that the critical role of NF-κB/IRF1 signaling in intra-tumoral cDC1 maturation may be conserved in human cancer and play an important role in disease progression. These findings could lead to new diagnostic and therapeutic approaches based on the NF-κB/IRF1 axis in cancer patients to improve immunotherapy and clinical outcomes.

## Material and Methods

### Mice

Mice were housed at Centre d’immunologie Marseille Luminy with water and food ad libitum and 12-h/12-h night/ daylight cycle under specific pathogen free conditions and handled in accordance with French and European directives. All animal experimentation was approved by Direction Départementale des Services Vétérinaires des Bouches du Rhône. C57BL/6J mice purchased from Janvier Labs. Experiments were performed with sex-matched littermate mice at 6–12 weeks of age. Xcr1Cre-mTFP1 (Xcr1-iCre in short) mice were previously described (Wohn et al.).

### Tumor cell cultures and injections

Mycoplasma free YUMM1.7 (*21*) or YUMMER1.7 (*22*) were cultured 37°C, 5% CO_2_ in RPMI1640 medium with 10% heat-inactivated fetal calf serum (Biochrom, Cambridge, UK), penicillin/streptomycin, and 50mM β-mercaptoethanol. YUMM1.7 or YUMMER1.7 cultures were washed in PBS pH 7.4 and harvested with PBS containing 2 mM EDTA, for 2 min at 37°C, then washed again in PBS. 1×10^6^ cells were injected s.c. in 100μl endotoxin-free PBS on the right flank of recipient mice. Tumor growth was measured using a digital caliper. Tumor volume stated in the figures refers to the Lxl^2^ considering the longest diameter (L) and its perpendicular (l) for each tumor.

### Tissue processing for flow cytometry

Tumors were minced and digested in RPMI 1640, 1 mg/ml Collagenase II (Sigma-Aldrich), 50 μg/ml DNaseI (Roche), and 0.1% (wt/vol) BSA for 30 min at 37°C with 750 rpm agitation. Cell suspensions were subsequently passed through a 70-μm cell strainer (BD Biosciences) and collected by centrifugation.

### Flow Cytometry

Flow cytometric analyses and cell sorting were performed using LSR Fortessa X20 or LSR-2 flow cytometers and Aria Cell sorter, respectively. Cells were pre-incubated with 2.4G2 antibody to block unspecific binding to Fc-receptors in all stainings. Staining with antibodies (see Antibodies in Key Resources Table) was performed in PBS, 2% FBS, 2 mM EDTA for 20 min at 4 °C. For exclusion of dead cells Live/Dead fixable cell stain kit (Invitrogen) was used in all experiments. HY-specific tetramers (Smcy_(738-746)_/H-2Db and Uty_(243-254)_/H-2Db) were provided from NIH Tetramer Core Facility and coupled to streptavidin-APC. Incubation with HY-tetramers was carried out before staining with antibodies at 10 nM for 40 min at 4 °C. Data were analysed using FlowJo (Tree Star, Inc.).

### Bone marrow derived DCs

Bone marrow derived DCs were generated as described (Mayer et al., 2014). Briefly, BM cells were incubated with 10 ng/ml GM-CSF (Peprotech, 315-03) and 1/10 FLT3L supernatant in RPMI1640 medium with 10% heat-inactivated fetal calf serum (Biochrom, Cambridge, UK), penicillin/streptomycin, and 50mM β-mercaptoethanol. BM cells were incubated in 10 ml complete medium with FLT3L combined with GM-CSF for 9 days and subsequently replated with the same combination of cytokines and harvested at day 15. Treatment with tumor conditioned medium (TCM) was carried out the last day before harvesting. TCM was produced by mincing YUMM1.7 tumor explants and incubating them at 50 mg/ml of RPMI1640 medium with 10% heat-inactivated fetal calf serum, penicillin/streptomycin, and 50mM β-mercaptoethanol for 24 hours at 37°C.

### Immunofluorescence imaging

Tumor pieces were fixed in 4% PFA for 4 h, washed in PBS pH7.4 for 4h, and dehydrated overnight in 30% sucrose in 0.1M PBS pH 7.4. Tissues were snap frozen, and 10 μm cryostat sections were stained with DAPI, anti-CD8b.2 (clone: 53-5.8) and anti-CD45 (clone 30-F11). Immunofluorescence confocal microscopy was performed by using a Zeiss LSM 780 confocal microscope using spectral unmixing with a 20× objective. Mozaic of 25 images were acquired for subsequent analysis. The number of CD8b^+^ cells was determined computationally by using a macroprogram developed for Fiji software encompassing plugins. Threshold and watershed plugins were used to calculate the algorithm allowing distinction of cells from signal background and separation of overlapping cells, respectively. The particle analysis plugin was set to calculate the number of cells whose range are from 55 to 65 μm^2^. The area of each tumor section was measured by setting masks on Dapi signal and using the toolset of measurement in Fiji on the inverted selected mask.

### Western blot analysis

Cells were washed and harvested in ice-cold PBS and pellets were lysed on ice in RIPA buffer (150 mm NaCl, 1% Nonidet P-40, 0.5% sodium deoxycholate, 0.1% SDS, 50 mm Tris, pH 8.0) containing a protease inhibitors cocktail (Sigma, P8340), 1 mM PMSF (1mM), 2mM PNPP, 10mM glycerol, 100 μM orthovanadate and 1 mM DTT. After 1 h of incubation on ice with frequent agitations, cell lysates were centrifuged at 12000 g, 10 min, the supernatants were collected and the concentration of proteins was determined using the DC Protein Assay, according to the manufacturer’s instructions (Bio-Rad Laboratories). Proteins (75 μg) from the various lysates were separated on 10-13% polyacrylamide slab gels (depending on the size of the protein to be analysed) and transferred to polyvinylidene fluoride membranes. The membranes were blocked with 5% skimmed milk in PBS for 1 h at room temperature and reacted for 16 h at 4°C with the appropriate primary antibody. Primary and HRP-conjugated antibodies were applied in 3% BSA in PBS, containing 0.02% sodium azide. Incubations with secondary antibodies were for 1 h at room temperature. Membranes were rinsed between incubations three times with PBS plus 0.05% tween-20. After the last wash, membranes were imaged using Amersham ECL prime (GE Healthcare, RPN2232).

### High-throughput gene expression analysis

Purification of total RNA from sorted cells was performed by using the RNeasy Plus Micro Kit (Qiagen) and concentration was determined using the Quant-IT RiboGreen RNA assay kit (Thermo Fisher Scientific). 2.5 ng total RNA was employed for first-strand cDNA synthesis with High Capacity cDNA Reverse Transcription Kit (Thermo Fisher Scientific) followed by a 15 cycles preamplification PCR of genes of interest using the Fluidigm PreAmp Master Mix (Fluidigm Europe B.V.) in accordance with the manufacturer’s instructions. Samples were subsequently treated with exonuclease I (New England Biolabs) to remove unincorporated primers and diluted 1:5 in Tris-EDTA buffer. High-throughput gene expression analysis was performed using the 96.96 or 48.48 dynamic arrays and Biomark HD system from Fluidigm in accordance with the manufacturer’s instructions and standard settings. Exon-spanning primers to amply genes of interest were designed using Primer-Blast (see Oligonucleotides in Key Resources Table). Obtained data were analysed using the Real-Time PCR Analysis Software (Fluidigm), and resulting CT values were normalized to *actin* or *ppia.* Heatmaps, Z-scores, and hierarchical clustering using the one minus Pearson correlation were generated using Morpheus (https://software.broadinstitute.org/morpheus/).

### Enrichment analyses

Gene set enrichment analysis (GSEA) on bulk RNA-seq data was run to calculate normalized enrichment score (NES) using metric for ranking genes: Diff_of_classes and classic enrichment statistics.

Enrichment analysis of scRNA-seq data was performed using the EnrichR tool (https://amp.pharm.mssm.edu/Enrichr/)21. The combined score is calculated by multiplying the log of the p-value computed with the Fisher exact test by the z-score computed by the correction to this test20. DEGs were selected upon setting an arbitrary cut-off for significance at p<0.00001 and adjusted pq 0.25. For the EnrichR analysis of the cluster markers from the WT dataset (Fig.2), only positive markers with an adj.p-val<0.05 were considered.

### Single-cell sorting and library preparation

Single cells were FACS-sorted into ice-cold 96-well PCR plates (Thermofisher) containing 2 μl lysis mix per well. The lysis mix contained 0.5 μl 0.4% (v/v) Triton X-100 (Sigma-Aldrich), 0.05 μl 40 U/μl RnaseOUT (Thermofisher), 0.08 μl 25 mM dNTP mix (Thermofisher), 0.5 μl 10 μM (dT)30_Smarter primer(*30*), 0.05 μl 0.5 pg/μl External RNA Controls Consortium (ERCC) spike-ins mix (Thermofisher), and 0.82 μl PCR-grade H20 (Qiagen). The plates containing single cells in lysis mix were stored at −80°C until further processing where they were thawed on ice, briefly spun down and incubated in a thermal cycler for 3 min at 72°C (lid temperature 72°C). Immediately after, the plate was placed back on ice and 3 μl RT mastermix was added to each well. The RT mastermix contained 0.25 μl 200 U/μl SuperScript II (Thermofisher), 0.25 μl 40 U/μl RnaseOUT (Thermofisher), and 2.5 μl 2× RT mastermix. The 2× RT mastermix contained 1 μl 5× SuperScript II buffer (Thermofisher), 0.25 μl 100 mM DTT (Thermofisher), 1 μl 5 M betaine (Sigma-Aldrich), 0.03 μl 1 M MgCl2 (Sigma-Aldrich), 0.125 μl 100 μM well-specific template switching oligonucleotide TSO_BCx_UMI5_TATA (Tables S3, S4), and 0.095 μl PCR-grade H2O (Qiagen). Reverse transcription was performed in a thermal cycler (lid temperature 70°C) for 90 min at 42°C, followed by 10 cycles of 2 min at 50°C and 2 min at 42°C, and then 15 min at 70°C. The plates with single-cell cDNA were stored at −20°C until further processing. For cDNA amplification, 7.5 μl LD-PCR mastermix was added to each well. The LD-PCR mastermix contained 6.25 μl 2× KAPA HiFi HotStart ReadyMix (Roche Diagnostics), 0.125 μl 20 μM PCR_Satija forward primer(*30*), 0.125 μl 20 μM SmarterR reverse primer (*30*) and 1 μl PCR-grade H2O (Qiagen). The amplification was performed in a thermal cycler (lid temperature 98°C) for 3 min at 98°C, followed by 22 cycles of 15 s at 98°C, 20 s at 67°C, 6 min at 72°C, and then a final elongation for 5 min at 72°C. The plates with amplified single-cell cDNA were stored at −20°C until further processing.

For library preparation, 5 μl of amplified cDNA from each well of a 96-well plate were pooled and completed to 500 μl with PCR-grade H2O (Qiagen). Two rounds of 0.6X solid-phase reversible immobilization beads (AmpureXP, Beckman, or CleanNGS, Proteigene) cleaning were used to purify 100 μl of pooled cDNA with final elution in 15 μl PCR-grade H2O (Qiagen). After quantification with Qubit dsDNA HS assay (Thermofisher), 800 pg purified cDNA pool was processed with the Nextera XT DNA sample Preparation kit (Illumina), according to the manufacturer’s instructions with modifications to enrich the 5’-ends of tagmented cDNA during library PCR. After tagmentation and neutralization, 25 μl tagmented cDNA was amplified with 15 μl Nextera PCR Mastermix, 5 μl Nextera i5 primer (S5xx, Illumina), and 5 μl of a custom i7 primer mix (0.5 μM i7_BCx + 10 μM i7_primer) (*30*). The amplification was performed in a thermal cycler (lid temperature 72°C) for 3 min at 72°C, 30 s at 95°C, followed by 12 cycles of 10 s at 95°C, 30 s at 55°C, 30 s at 72°C, and then a final elongation for 5 min at 72°C. The resulting library was purified with 0.8X solid-phase reversible immobilization beads (AmpureXP, Beckman, or CleanNGS, Proteigene). Libraries generated from multiple 96-well plates of single cells and carrying distinct i7 barcodes were pooled for sequencing on an Illumina NextSeq550 platform, with High Output 75 cycles flow cells, targeting 5 × 105 reads per cell in paired-end single-index mode with the primers previously reported (*30*) and cycles: Read1 (Read1_SP, 67 cycles), Read i7 (i7_SP, 8 cycles), and Read2 (Read2_SP, 16 cycles).

### Single-cell RNA sequencing data analysis FB5P-seq

Raw fastq files were processed to generate single-cell unique molecular identifier (UMI) count matrices as described (PMID: 32194545). Read alignment on the mouse reference genome (GRCm38.94 with ERCC92) was performed using STAR (v2.5.3a). Digital gene expression data were extracted using Drop-seq software (v1.12).

The counts matrix was loaded to R (3.6.1) and Seurat (v3.1.5) (PMID: 29608179) was used for downstream analyses in a similar way as in Abbas et al. 2020 (PMID: 32690951). First, we selected the genes that were expressed in at least 5 cells. Then, we excluded the cells that had less than 600 detected genes or more than 10% mitochondrial genes.

We identified contaminants based on two approaches i) performing connectivity map analysis (cmap) using homemade signatures ii) using FindMarkers functions in seurat to identify the genes for characterizing the clusters. These two approached allowed us to identify and remove five clusters (C4,C9,C10,C12 and C13, n=110) as contaminants.

### Analysis of scRNA-seq data of Figure 2 and Supplementary Figure 2

We obtained 113 bona fide cDC1. Gene expression is shown as log(normalized values) and protein expression as inverse hyperbolic arcsine (asinh) of fluorescence intensity. Afterwards, PCA was performed using the highly variable genes. A Shared Nearest Neighbor (SNN) Graph was built using the 4 first principal components with the FindNeighbors function (dims = 1:4, k.param=20). Clustering was then performed with a resolution of 1.3. For dimensionality reduction, we performed a UMAP using the RunUMAP function (n.neighbors=7, spread=1, min.dist=2.5).

### Analysis of scRNA-seq data of Figure 6 and Supplementary Figure 6

We obtained 360 bona fide cDC1. A SNN Graph was built using the 6 first principal components. Clustering was performed with a resolution of 0.6. A UMAP was performed using the RunUMAP function (n.neighbors=7, spread=1, min.dist=0.6). The differentially expressed genes were determined using the FindMarkers function (test.use= “bimod”, min.pct= 0.5).

### Heatmaps

Heatmaps based on Z-scores and hierarchical clustering using the one minus Pearson correlation were generated using Morpheus (https://software.broadinstitute.org/morpheus/).

### Analysis of scRNA-seq data from lung adenocarcinoma

ScRNA-seq data of DCs from the lungs of wild-type tumor-naïve and tumor-bearing mice were downloaded from GEO using accession code GSM3832735 and GSM3832737 (*15*). The count matrix was loaded to R and Seurat as described above. We used the same parameters and the same strategy as described above to exclude contaminating cells. From 4695 starting cells, we retained 1418 cells identified as cDC1.

### Alignment of clusters from melanoma and lung adenocarcinoma datasets

Marker genes of clusters for each dataset were extracted separately, using the FindMarkers function of Seurat and the “bimod” parameter. We then computed the Jaccard Indices (JI) between each cluster of one dataset with any other cluster of the other dataset, as already published(*51*). The JI measures similarity between finite sample sets, and is defined as the size of the intersection divided by the size of the union of the sample sets: J(A,B)= |A ⋂ B| / |A U B|. The JI matrix was then used to generate a heatmap.

### BubbleMap

High-throughput GSEA was performed to assess the enrichment of selected transcriptomic signatures across clusters of the scRNA-seq data, using BubbleMap from the BubbleGUM (v1.3.19) suite(*52*), with the default parameters except: Number of permutations: 6000, Max geneset size= 5000, Min geneset size= 10. We used the Hallmark and Kegg geneset collections (v7.0) from the MSigDB(*53*). We also used genesets of genes upregulated (Mat ON/Core UP) and downregulated (Mat OFF/Core DN) during DC maturation (*31, 32*).

### Pseudotemporal analysis

We used Monocle (v3) to investigate cell trajectories and the dynamics of gene expression within these trajectories. The input cells and genes were selected using Seurat. We first clustered and projected the cells onto a low-dimensional space (UMAP) calculated by Seurat. Finally, Monocle resolved the activation trajectories and calculated along them the pseudo-time for each cell using monocle functions reduce_dimensions(max_components=3; reduction_method=“UMAP”; preprocess_method = “PCA”; umap.min_dist = 0.8; umap.n_neighbors = 10) and cluster_cells(resolution= 2e-2; reduction_method = “UMAP”; k = 15; num_iter = 2;).

### Computational analysis of cancer patient data

Datasets for all the melanoma patients whose tumors were profiled with RNA-seq were downloaded from the GDC Data Portal (https://portal.gdc.cancer.gov/). This resulted in a cohort of 468 patients, all coming from the The Cancer Genome Atlas (TCGA)-SKCM project. The expression values, FPKM (fragments per kilobase of exon model per million reads mapped), for each of the selected genes across these patients were retrieved and Gene Signature (GS) scores calculated for each of them. These scores are defined as the average expression of the genes forming the defined GS. Subcohorts were identified using a given GS (e.g. patients with cDC1-enriched tumors or those in top quartile of a given GS). The annotated survivals of all the patients in the identified subcohorts were retrieved from the GDC Data Portal to draw the corresponding Kaplan-Meier survival plots.

Heatmaps, Z-scores, and hierarchical clustering using the one minus Pearson correlation were generated using Morpheus (https://software.broadinstitute.org/morpheus/). Gene signature analyses for BRCA database from TCGA was generated using GEPIA2 (http://gepia2.cancer-pku.cn/) (*54*).

### Statistical analysis

When referred by stars, *p* values were calculated using either the Mann Whitney test or the Fisher’s exact test when appropriate using GraphPad prism software. *p < 0.05, **p < 0.01, ***p < 0.001 and ****p < 0.0001.

## Acknowledgements

These studies were supported institutional funding from the French National Institute of Health and Medical Research (INSERM), Centre national de la recherche scientifique (CNRS), Aix-Marseille-Universite (AMU), and by grants to T.L. from the French Ligue Nationale contre le Cancer (LNCC Equipe Labellisee EL2016-3) and Institut National du Cancer (INCa PLBIO13-244), to K.C. from ARC (PJA20181207726) and to M.D. from INCa (PLBIO 2018-152). GG was funded by LNCC and Marie-Curie actions IEF-Horizon2020 under REA grant agreement no. 702933. ASC was funded by a PhD fellowship from the Turing Center for Living Systems (CENTURI).

We thank Anna Baranska and Noudjoud Attaf for operational help in flow cytometry and sample preparation for scRNAseq, respectively. We thank the CIML bioinformatics, cytometry and histology platform for their technical and methodological help.

## Author contributions

GG performed most of the experiments and analyses and wrote the manuscript. TPVM supervised scRNA-seq analyses and performed BubbleGum analyses. ASC performed scRNA-seq data analyses. EB performed experiments characterizing immunogenicity of YUMM1.7 models. CV maintained transgenic mouse strains. KC generated *Xcr1^DTA^* mice. NAA established YUMM1.7 models and their analyses and supervised YUMM1.7 immunogenicity experiments. PJB performed computational analyses of TCGA datasets. AN assisted with FB5P-seq library preparation. CD performed primary bioinformatics analysis of FB5P-seq data. PM provided technical support and advice for FB5P-seq strategy. BM provided access to *Xcr1-iCre* mice prior to their publication. MD supervised the studies and contributed to manuscript writing. TL conceived and supervised the studies, provided funding support and contributed to manuscript writing.

