## Supplementary figures and legends for "An NF-κB/IRF1 axis programs cDC1s to drive anti-tumor immunity"

#### Supplementary Figure1

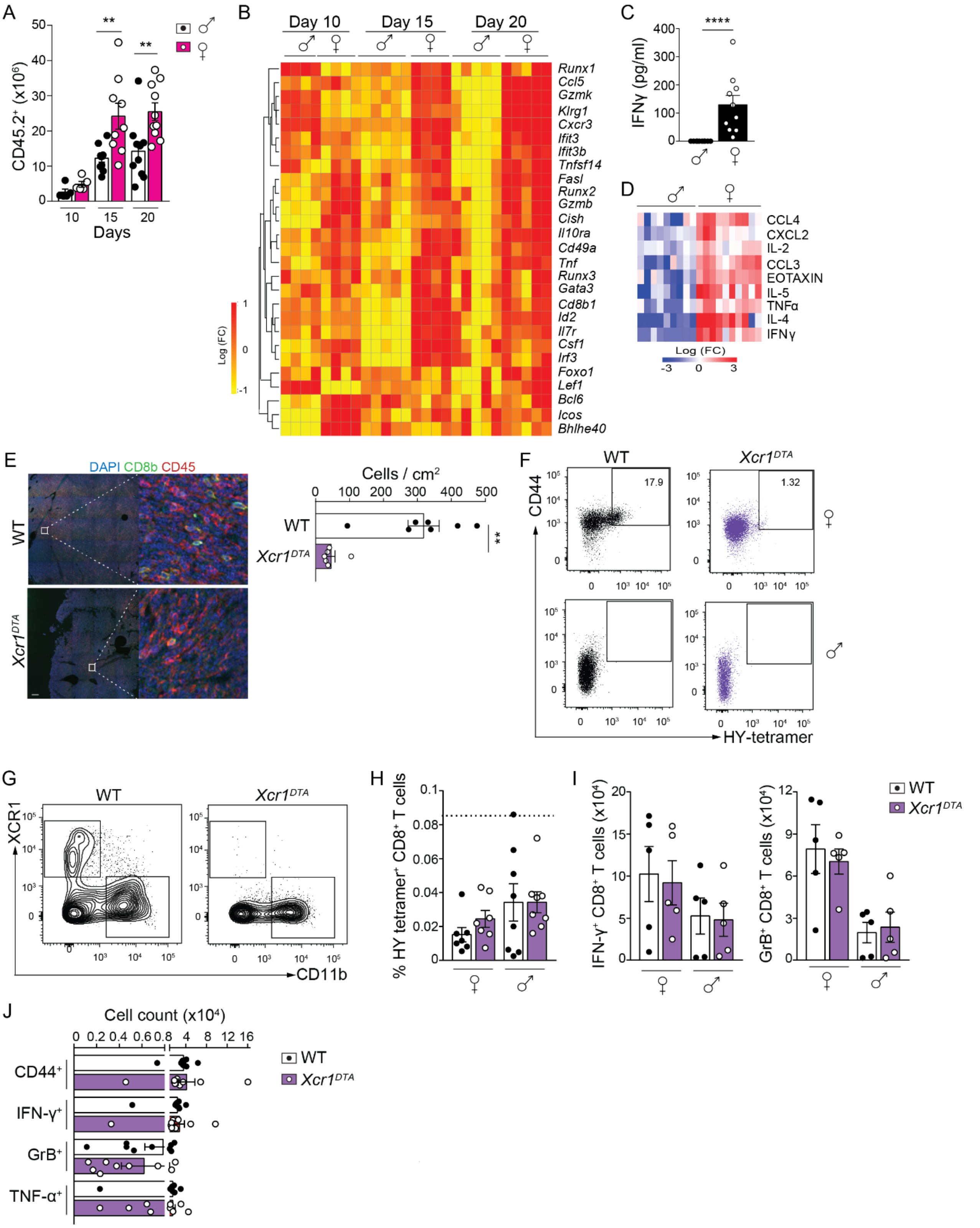

**Figure S1. cDC1s are required for control of immunogenic tumor growth.**

(**A**) Total numbers of CD45.2$^{+}$ cells per 250mg of tumor tissue in YUMM1.7 tumor-bearing male and female mice.

(**B**) Differential expression of selected genes involved in the regulation of T cell activation by CD8$^{+}$ T cells in YUMM1.7 tumors from male and female mice.

(**C**,**D**) Analysis cytokines in YUMM1.7 tumour lysates from male and female mice at 10 days; (**C**) IFN-$\gamma$ ELISA and (**D**) multiplex Luminex assay.

(**E**) Confocal microscopy of CD8$^{+}$ T cells in sections of YUMM1.7 tumors from WT or *Xcr1*$^{DTA}$ female mice (scale bar 200 µM; zoom x3000 magnification).

(**F**) Representative flow cytometry plots of HY-tetramer$^{+}$ CD8$^{+}$ T cells in YUMM1.7 tumors from WT or *Xcr1*$^{DTA}$ male and female mice.

(**G**) Representative flow cytometry plots for analysis of cDC1 (XCR1$^{+}$) and cDC2 (CD11b$^{+}$) in TDLNs from WT and *Xcr1*$^{DTA}$ female mice bearing YUMM1.7 tumors. DCs are gated as CD11c$^{+}$ MHCII$^{+}$ Lin$^{-}$ (CD3$^{-}$ CD19$^{-}$ NK1.1$^{-}$ Ly6G$^{-}$) cells.

(**H**) Flow cytometry analysis of HY-tetramer$^{+}$ CD8$^{+}$ T cells in TDLNs from WT and *Xcr1*$^{DTA}$ male or female mice engrafted with YUMM1.7 tumors. The dashed line represents the average staining obtained with control HPV16 tetramers.

(**I**) Numbers of activated IFN-$\gamma$ and GrB-expressing CD8$^{+}$ T cells in TDLNs from WT and Xcr1$^{DTA}$ male or female mice engrafted with YUMM1.7 tumors.

(**J**) Numbers of activated CD8$^{+}$ T cells expressing CD44, IFN-$\gamma$ or GrB in non-draining lymph nodes from WT and Xcr1$^{DTA}$ male mice engrafted with YUMMER1.7 tumors.

**Supplementary Figure 2**

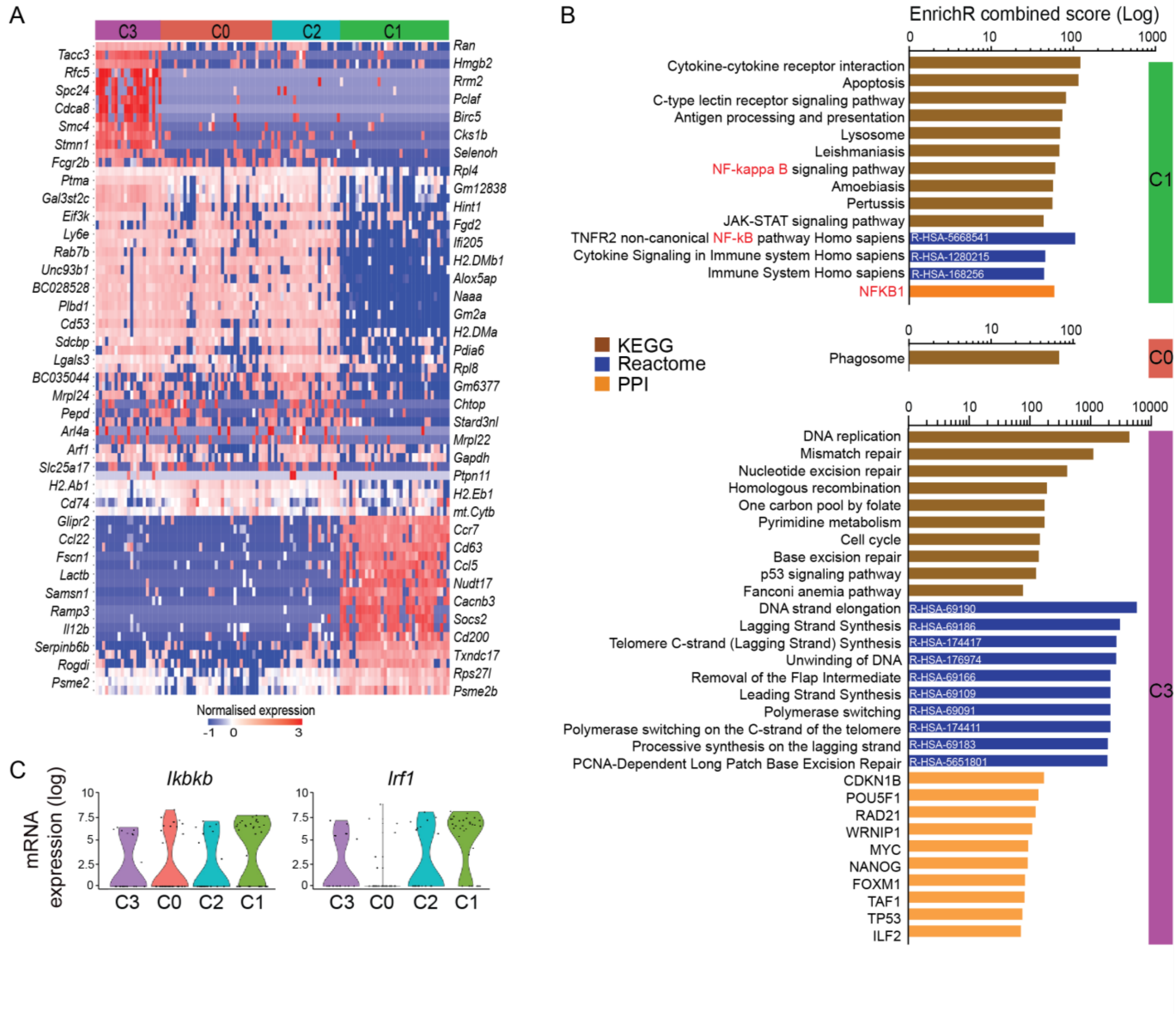


**Figure S2. Single-cell transcriptional profiling of intra-tumoral cDC1s.**

(**A**) Gene expression heatmap of the top 20 markers defining Seurat clusters in Figure 2A (C0-3).

(**B**) Annotation enrichment associated with the positive markers (adj. p-val<0.05) of each cluster performed using the EnrichR package, against selected gene annotation databases (KEGG, Reactome and Protein-Protein Interaction (PPI)). No significant enrichment was observed for C2.

(**C**) Violin plots showing the expression of *Ikbkb* and *Irf1* across clusters C0-3.

**Supplementary Figure 3**

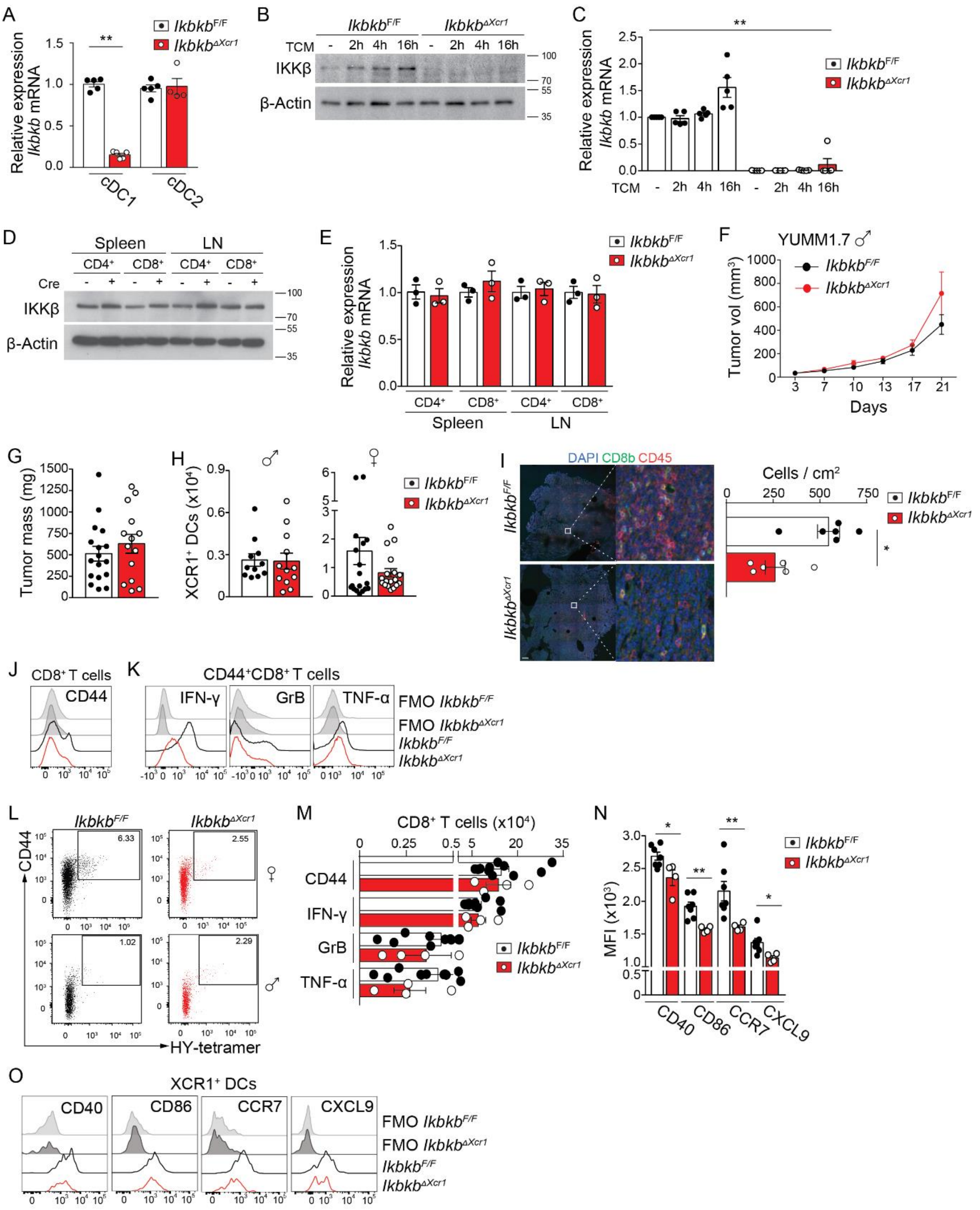

**Figure S3. Conditional deletion of IKKβ in cDC1s.**

(**A**) *Ikbkb* mRNA expression analyzed by qRT-PCR in cDC1s and cDC2s isolated by flow cytometry from pooled spleens and LNs, expression was normalized to *Actb* mRNA.

(**B**,**C**) Bone marrow-derived DCs (BMDCs) from $Ikbkb^{F/F}$ and $Ikbkb^{\Delta Xcr1}$ mice were treated with YUMM1.7 tumor-conditional medium (TCM) for the indicated times. cDC1s were sorted by flow cytometry and (**B**) protein extracts prepared for western blot or (**C**) RNA isolated for qRT-PCR analysis of Ikbkb expression.

(**D**,**E**) $CD4^+$ and $CD8^+$ T cells were sorted by flow cytometry from spleen and LNs of $Ikbkb^{F/F}$ and $Ikbkb^{\Delta Xcr1}$ mice and analyzed by (**D**) western blotting or (**E**) qRT-PCR.

(**F**) YUMM1.7 tumor growth in $Ikbkb^{F/F}$ (n=10) and $Ikbkb^{\Delta Xcr1}$ (n=8) male mice, data is represented as mean ± SEM.

(**G**) Comparison of tumour burden between $Ikbkb^{F/F}$ and $Ikbkb^{\Delta Xcr1}$ male mice engrafted with YUMM1.7 tumors.

(**H**) Numbers of cDC1s per 250mg of tumor tissue in $Ikbkb^{F/F}$ and $Ikbkb^{\Delta Xcr1}$ male and female mice bearing YUMM1.7 tumors.

(**I**) Confocal microscopy of $CD8^+$ T cells in sections of YUMM1.7 tumors from $Ikbkb^{F/F}$ and $Ikbkb^{\Delta Xcr1}$ female mice (scale bar 200 μM; zoom x3000 magnification).

(**J**,**K**) Representative histograms from flow cytometry analysis of activated (**J**) $CD44^+$ $CD8^+$ T cells and (**K**) IFN-γ, GrB or TNF-α expression in YUMM1.7 tumors from female $Ikbkb^{F/F}$ and $Ikbkb^{\Delta Xcr1}$ mice. FMO controls are shown.

(**L**) Representative flow cytometry plots of HY-tetramer$^+$ $CD8^+$ T cells in YUMM1.7 tumors from $Ikbkb^{F/F}$ and $Ikbkb^{\Delta Xcr1}$ male and female mice.

(**M**) Numbers of activated $CD8^+$ T cells expressing CD44, IFN-γ or GrB in TDLNs from $Ikbkb^{F/F}$ and $Ikbkb^{\Delta Xcr1}$ male mice engrafted with YUMMER1.7 tumors.

(**N**) Expression of maruration markers (CD40, CD86, CCR7) and CXCL9 by migratory cDC1s in TDLNs from $Ikbkb^{F/F}$ and $Ikbkb^{\Delta Xcr1}$ male mice bearing YUMMER1.7 tumors assessed by flow cytometry and expressed as mean fluorescence intensity (MFI). (**O**) Representative histograms from flow cytometry analysis in (**N**). FMO controls are shown.

### Supplementary Figure 4

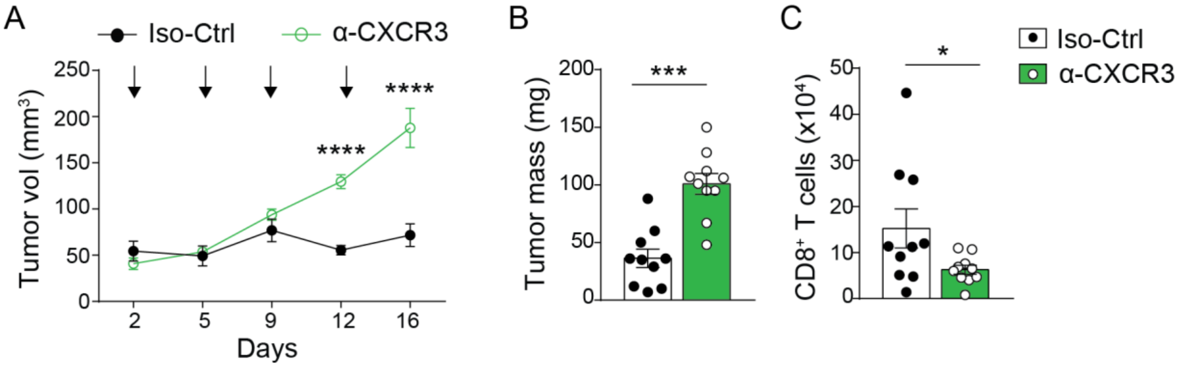


**Figure S4. CXCR3 blockade increases growth of immunogenic tumors.**

(**A**,**B**) YUMM1.7 tumor growth in female mice treated with anti-CXCR3 (300μg) or control antibody (Iso-Ctrl) at indicated times (arrows). Data are shown as mean ± SEM, n=10 in each group, and are representative of two independent experiments.

(**C**) Numbers of $CD8^+$ T cells per 250mg of tumor tissue in mice treated anti-CXCR3 treated mice. Data are shown as mean ± SEM; each point corresponds to one mouse.

Statistical analysis was performed with two-way ANOVA followed by Sidak's multiple comparison tests (**A**) and Mann-Whitney test (**B**,**C**).

### Supplementary Figure 5

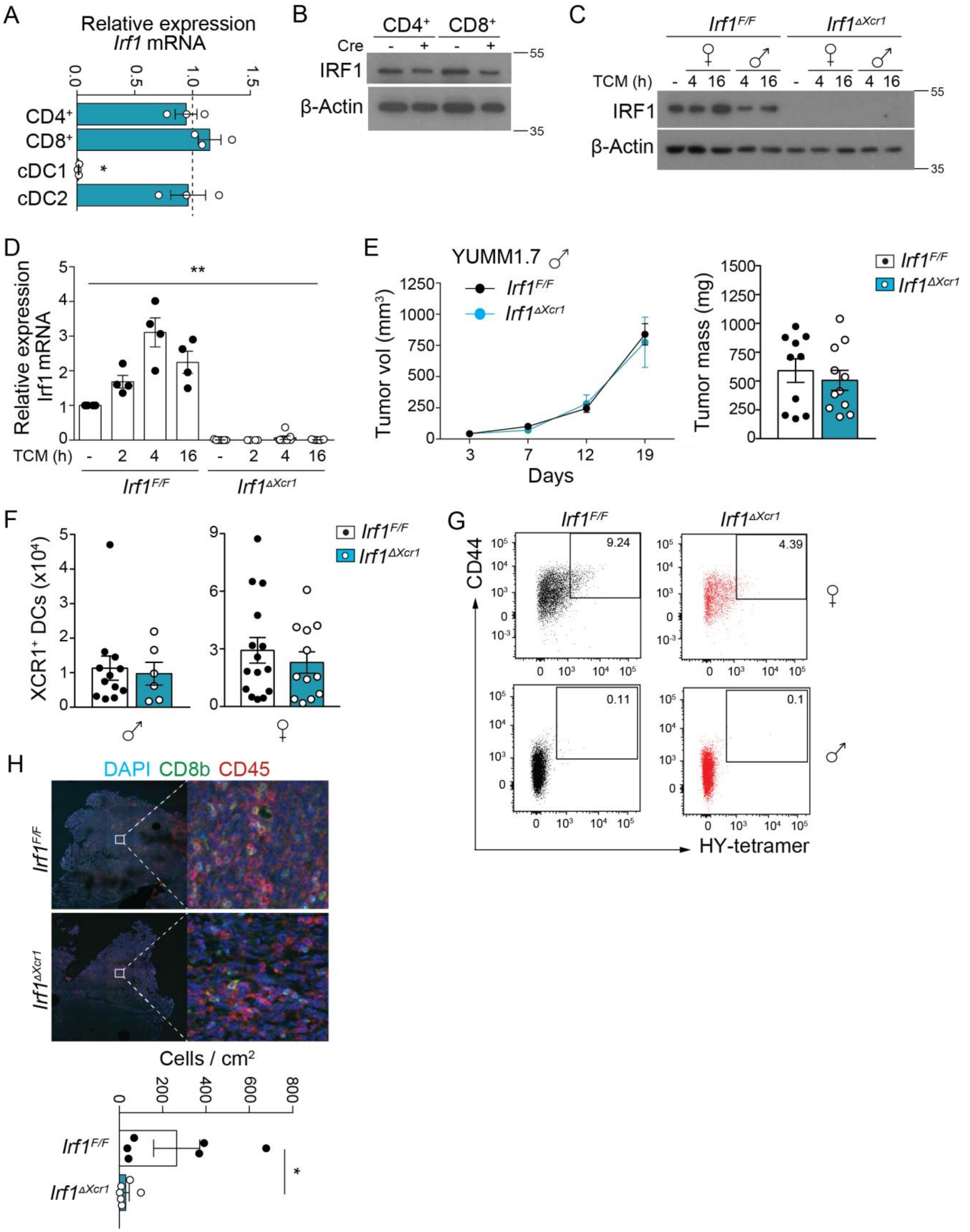

**Figure S5. Conditional deletion of IRF1 in cDC1s.**

(**A**) *Irf1* mRNA expression analyzed by qRT-PCR in $CD4^+$ and $CD8^+$ T cells, cDC1s and cDC2s isolated by flow cytometry from pooled spleens and LNs of $Irf1^{F/F}$ and $Irf1^{\Delta Xcr1}$ mice, expression was normalized to *Actb* mRNA. Data represents relative expression normalized to $Irf1^{F/F}$ mice.

(**B**) Western blot analysis of Irf1 expression in sorted $CD4^+$ and $CD8^+$ T cells.

(**C**,**D**) BMDCs from $Ikbkb^{F/F}$ and $Ikbkb^{\Delta Xcr1}$ mice were treated with TCM for the indicated times, cDC1s were sorted by flow cytometry and (**C**) protein extracts prepared for western blot or (**D**) RNA isolated for qRT-PCR analysis of Irf1 expression.

(**E**) YUMM1.7 tumor growth in $Irf1^{F/F}$ (n= 9) and $Irf1^{\Delta Xcr1}$ (n=5) male mice, data is represented as mean ± SEM.

(**F**) Numbers of cDC1s per 250mg of tumor tissue in $Irf1^{F/F}$ and $Irf1^{\Delta Xcr1}$ male and female mice bearing YUMM1.7 tumors.

(**G**) Representative flow cytometry plots of HY-tetramer$^+$ $CD8^+$ T cells in YUMM1.7 tumors from $Irf1^{F/F}$ and $Irf1^{\Delta Xcr1}$ male and female mice.

(**H**) Confocal microscopy of $CD8^+$ T cells in sections of YUMM1.7 tumors from $Ikbkb^{F/F}$ and $Ikbkb^{\Delta Xcr1}$ female mice (scale bar 200 µM; zoom x3000 magnification).

**Supplementary Figure 6**

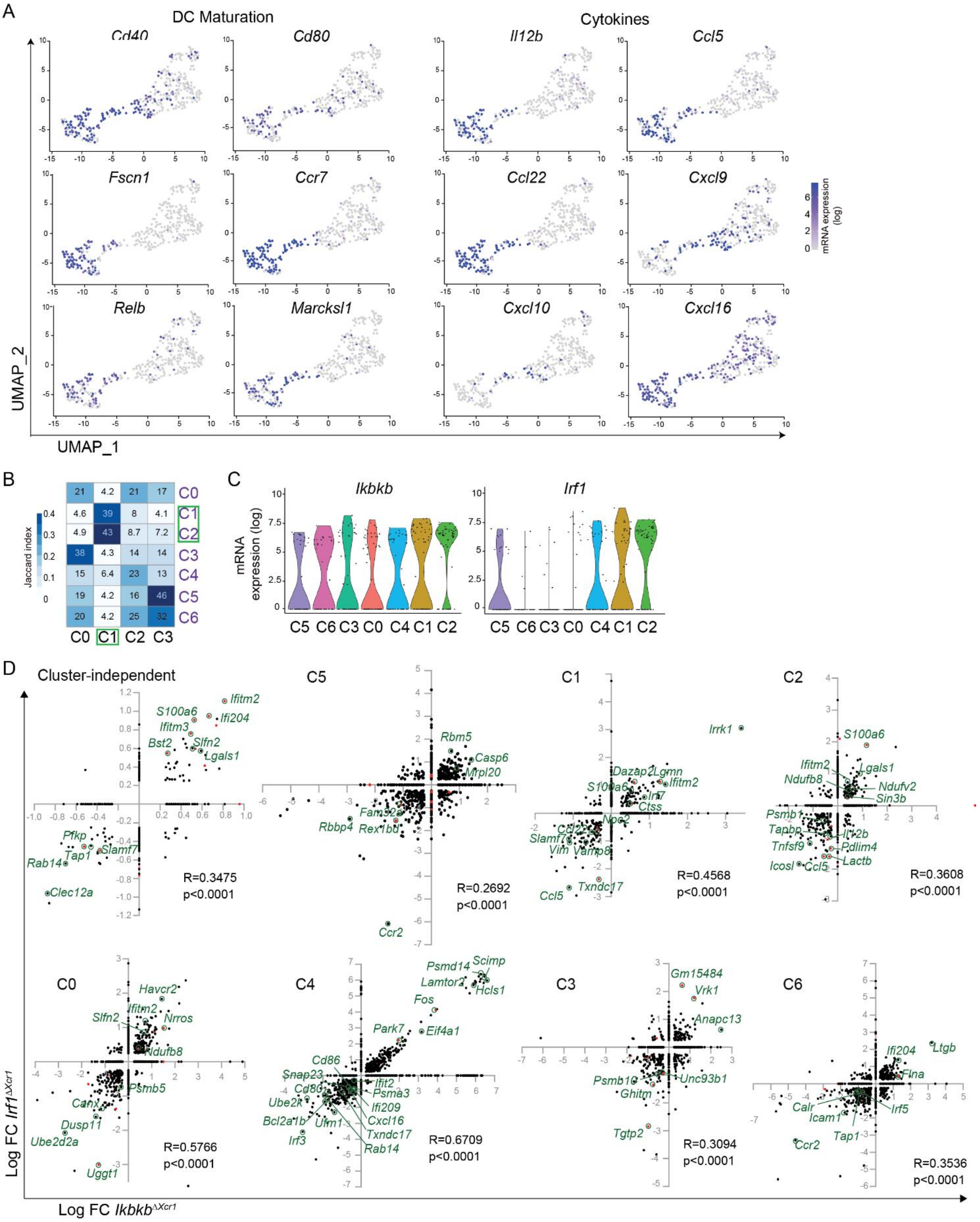

### Supplementary Figure 6 (cont.)

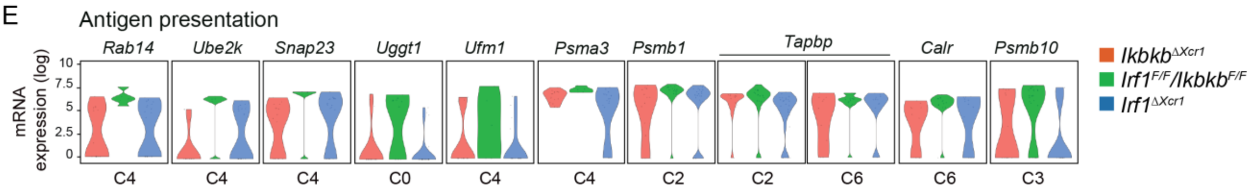


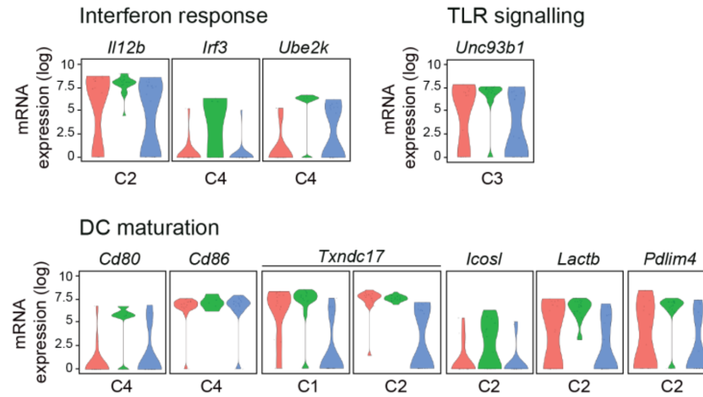


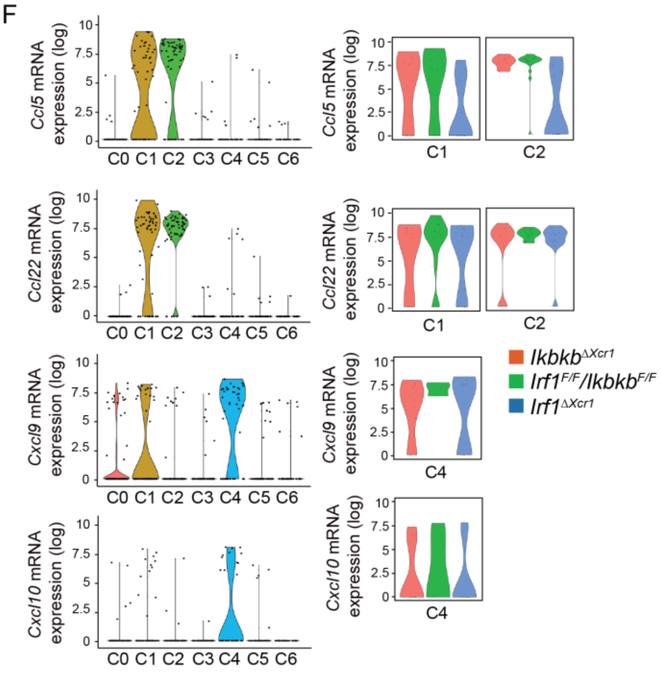


G

#### DEGs in *Ikbkb*$^{\Delta Xcr1}$

| C | Genes | logFC | *p* | adjusted *p* |
|---|---|---|---|---|
| C-ind | *Aif1* | 0.47 | $5.12E^{-8}$ | 0.0059 |
| C-ind | *Sept7* | -0.36 | $4.67E^{-7}$ | 0.0054 |
| C-ind | *Irf5* | -0.46 | $3.16E^{-6}$ | 0.0036 |
| C-ind | *Ifitm2* | 0.81 | $5.52E^{-6}$ | 0.0638 |
| C-ind | *Slamf7* | -0.37 | $9.89E^{-6}$ | 0.1142 |
| C-ind | *Mrps18c* | 0.96 | $1.54E^{-5}$ | 0.1779 |
| C-ind | *Ubb* | -0.41 | $1.71E^{-5}$ | 0.1978 |
| C0 | *Eef1d* | 0.97 | $3.60E^{-7}$ | 0.0041 |
| C0 | *Ndufb10* | -0.43 | $6.56E^{-7}$ | 0.0076 |
| C0 | *Rab14* | -0.77 | $1.30E^{-6}$ | 0.0150 |
| C0 | *Slc8b1* | -1.71 | $2.04E^{-6}$ | 0.0236 |
| C0 | *Celf2* | -0.78 | $2.06E^{-6}$ | 0.0238 |
| C0 | *Nhp2* | -0.53 | $4.82E^{-6}$ | 0.0557 |
| C0 | *Selplg* | -0.51 | $8.50E^{-6}$ | 0.0982 |
| C0 | *Lsp1* | -0.32 | $1.29E^{-5}$ | 0.1494 |
| C0 | *Bub3* | 1.50 | $1.41E^{-5}$ | 0.1626 |
| C0 | *Ndufb8* | 0.42 | $1.64E^{-5}$ | 0.1895 |
| C0 | *Rps19* | 0.31 | $1.71E^{-5}$ | 0.1980 |
| C1 | *Npc2* | -0.36 | $7E^{-6}$ | 0.0809 |
| C2 | *Reep5* | 0.61 | $3.36E^{-7}$ | 0.0039 |
| C2 | *Psmb1* | -0.54 | $6.05E^{-7}$ | 0.0070 |
| C2 | *Sin3b* | 0.41 | $9.97E^{-7}$ | 0.0115 |
| C2 | *Il2rg* | 0.56 | $1.86E^{-6}$ | 0.0215 |
| C2 | *Pdlim4* | -0.28 | $3.17E^{-6}$ | 0.0366 |
| C2 | *Ly6e* | 5.56 | $8.85E^{-6}$ | 0.1022 |
| C2 | *Rer1* | -0.51 | $1.39E^{-5}$ | 0.1612 |
| C3 | *Rps14* | 0.28 | $2.36E^{-9}$ | $2.72E^{-5}$ |
| C3 | *Pp1ca* | -0.86 | $4.21E^{-9}$ | $4.86E^{-5}$ |
| C3 | *Vdac2* | -1.81 | $1.39E^{-8}$ | 0.0001 |
| C3 | *Anxa2* | -0.29 | $4.1E^{-8}$ | 0.0005 |
| C3 | *Stard3nl* | -1.15 | $7.99E^{-8}$ | 0.0009 |
| C3 | *Arpc1b* | -0.30 | $6.61E^{-7}$ | 0.0076 |
| C3 | *Ddx5* | -0.43 | $1.04E^{-6}$ | 0.0120 |
| C3 | *Hnrnpk* | -0.65 | $2.36E^{-6}$ | 0.0273 |
| C3 | *Naaa* | -0.28 | $3.92E^{-6}$ | 0.0454 |
| C3 | *Tpm4* | -0.88 | $4.61E^{-6}$ | 0.0533 |
| C3 | *Tmem59* | -0.32 | $4.76E^{-6}$ | 0.0550 |
| C3 | *Myl6* | 0.34 | $4.84E^{-6}$ | 0.0559 |
| C3 | *Sh3brgl3* | 0.37 | $9.43E^{-6}$ | 0.1089 |
| C3 | *Ghitm* | -0.74 | $1.66E^{-5}$ | 0.1924 |
| C3 | *Rab7b* | -0.58 | $1.88E^{-5}$ | 0.2171 |
| C4 | *Rpl14* | 0.47 | $7.64E^{-9}$ | $8.83E^{-5}$ |
| C4 | *Laptm5* | 0.43 | $3.37E^{-8}$ | 0.0004 |
| C4 | *Park7* | 1.97 | $2.42E^{-7}$ | 0.0028 |
| C4 | *Arf4* | 1.17 | $2.99E^{-7}$ | 0.0035 |
| C4 | *Spcs1* | 0.69 | $5.04E^{-7}$ | 0.0058 |
| C4 | *Rps3* | 0.28 | $7.75E^{-6}$ | 0.0895 |
| C4 | *Cox4i1* | 1.15 | $8.59E^{-6}$ | 0.0993 |
| C4 | *Ppia* | 0.61 | $1.11E^{-5}$ | 0.1282 |
| C4 | *Ccdc12* | 0.44 | $1.55E^{-5}$ | 0.1789 |
| C4 | *Fos* | 3.90 | $2.11E^{-5}$ | 0.2435 |
| C5 | *Hnrnpk* | -0.31 | $8.60E^{-9}$ | $9.94E^{-5}$ |
| C5 | *Ctsc* | -0.57 | $1.18E^{-7}$ | 0.0014 |
| C5 | *Mrpl20* | 0.88 | $4.39E^{-7}$ | 0.0051 |
| C5 | *Rex1bd* | -1.27 | $7.26E^{-7}$ | 0.0084 |
| C5 | *Fam32a* | -1.09 | $3.09E^{-6}$ | 0.0357 |
| C5 | *Mrpl35* | -2.18 | $3.58E^{-6}$ | 0.0414 |
| C5 | *Vdac3* | -0.81 | $4.33E^{-6}$ | 0.0499 |
| C5 | *Herpud1* | -1.16 | $1.03E^{-5}$ | 0.1188 |
| C5 | *Mtpn* | -0.78 | $1.62E^{-5}$ | 0.1869 |
| C5 | *Sh3bgrl* | 0.64 | $2.14E^{-5}$ | 0.2478 |
| C6 | *Tpt1* | -1.09 | $6.27E^{-6}$ | 0.0724 |
| C6 | *Tap1* | -0.75 | $6.56E^{-6}$ | 0.0758 |
| C6 | *Slbp* | -2.90 | $8.23E^{-6}$ | 0.0951 |
| C6 | *Ankrd12* | -2.15 | $1.66E^{-5}$ | 0.1916 |
| C6 | *Flna* | 1.28 | $2.05E^{-5}$ | 0.2366 |

#### DEGs in *Irf1*$^{\Delta Xcr1}$

| C | Genes | logFC | *p* | adjusted *p* |
|---|---|---|---|---|
| C-ind | *Ifitm2* | 1.11 | $8.08E^{-18}$ | $9.34E^{-14}$ |
| C-ind | *S100a6* | 0.91 | $7.50E^{-12}$ | $8.67E^{-8}$ |
| C-ind | *Fuca1* | -0.69 | $7.95E^{-9}$ | $8.77E^{-5}$ |
| C-ind | *Irf7* | 0.95 | $1.28E^{-8}$ | 0.0001 |
| C-ind | *Tmem258* | -0.42 | $2.11E^{-8}$ | 0.0002 |
| C-ind | *Pfdn5* | -0.29 | $3.24E^{-8}$ | 0.0003 |
| C-ind | *Zbp1* | 0.36 | $1.46E^{-7}$ | 0.0017 |
| C-ind | *Ifitm3* | 0.76 | $1.53E^{-7}$ | 0.0018 |
| C-ind | *Pfkp* | -0.46 | $1.81E^{-7}$ | 0.0076 |
| C-ind | *Plac8* | 0.85 | $1.98E^{-6}$ | 0.0021 |
| C-ind | *Ctsc* | 0.60 | $2.08E^{-6}$ | 0.0241 |
| C-ind | *Txn-ps1* | -0.76 | $2.10E^{-6}$ | 0.0248 |
| C-ind | *Ctsh* | 0.39 | $2.52E^{-6}$ | 0.0291 |
| C-ind | *Bst2* | 0.55 | $1.91E^{-5}$ | 0.2208 |
| C-ind | *Klrk1* | 0.41 | $2.04E^{-9}$ | 0.2354 |
| C0 | *Nrros* | 0.98 | $2.16E^{-9}$ | $2.50E^{-5}$ |
| C0 | *Dad1* | 0.87 | $3.83E^{-7}$ | 0.0044 |
| C0 | *Ssr4* | 0.32 | $1.24E^{-6}$ | 0.0143 |
| C0 | *Rps23* | 0.38 | $2.97E^{-6}$ | 0.0343 |
| C0 | *Uggt1* | -3.00 | $4.21E^{-6}$ | 0.0486 |
| C0 | *Cd53* | 0.42 | $9.23E^{-6}$ | 0.1066 |
| C0 | *Sbds* | 0.62 | $1.40E^{-5}$ | 0.1615 |
| C1 | *Lcp1* | 0.63 | $2.80E^{-8}$ | 0.0003 |
| C1 | *Lgmn* | 1.15 | $2.48E^{-7}$ | 0.0029 |
| C1 | *Ddx5* | 0.69 | $5.02E^{-7}$ | 0.0058 |
| C1 | *Dazap2* | 1.11 | $7.22E^{-7}$ | 0.0083 |
| C1 | *Stat1* | 1.12 | $7.34E^{-7}$ | 0.0085 |
| C1 | *Hnrnpk* | 0.31 | $1.91E^{-6}$ | 0.0220 |
| C1 | *Tpm3* | 0.47 | $2.22E^{-6}$ | 0.0257 |
| C1 | *Txndc17* | -2.37 | $3.21E^{-6}$ | 0.0371 |
| C1 | *Cd83* | 0.43 | $5.21E^{-6}$ | 0.0602 |
| C1 | *Rplp0* | -0.51 | $5.52E^{-6}$ | 0.0638 |
| C1 | *Npc2* | -0.61 | $1.21E^{-5}$ | 0.1394 |
| C1 | *Ctss* | 0.35 | $1.31E^{-5}$ | 0.1510 |
| C2 | *Txndc17* | -1.80 | $5.55E^{-11}$ | $6.41E^{-7}$ |
| C2 | *Pdlim4* | -1.35 | $1.37E^{-8}$ | 0.0001 |
| C2 | *S100a6* | 1.91 | $1.97E^{-8}$ | 0.0002 |
| C2 | *Ndufb8* | 0.48 | $4.57E^{-8}$ | 0.0005 |
| C2 | *Ctsc* | 1.28 | $5.11E^{-8}$ | 0.0006 |
| C2 | *Selenow* | -0.49 | $2.20E^{-7}$ | 0.0025 |
| C2 | *Sin3b* | 0.25 | $3.14E^{-7}$ | 0.0036 |
| C2 | *Gapdh* | 0.52 | $3.75E^{-7}$ | 0.0043 |
| C2 | *Slc25a4* | -0.66 | $3.37E^{-8}$ | 0.0044 |
| C2 | *Rad23a* | 0.82 | $4.95E^{-7}$ | 0.0057 |
| C2 | *Ccl5* | -1.41 | $7.67E^{-7}$ | 0.0089 |
| C2 | *H2-t23* | -0.87 | $1.03E^{-6}$ | 0.0119 |
| C2 | *Serpinb9* | -0.44 | $1.01E^{-5}$ | 0.1167 |
| C2 | *Lactb* | -1.60 | $1.27E^{-5}$ | 0.1470 |
| C2 | *Psme1* | -0.53 | $1.37E^{-5}$ | 0.1678 |
| C2 | *Idi1* | 1.16 | $1.45E^{-5}$ | 0.1626 |
| C2 | *Ndufv2* | 0.53 | $1.56E^{-5}$ | 0.1799 |
| C3 | *Unc93b1* | -0.95 | $2.41E^{-8}$ | 0.0003 |
| C3 | *Vrk1* | 1.77 | $1.49E^{-6}$ | 0.0173 |
| C3 | *Tgtp2* | -2.84 | $5.47E^{-6}$ | 0.0632 |
| C3 | *Cfl1* | 0.30 | $8.36E^{-6}$ | 0.0966 |
| C3 | *Gm15484* | 2.22 | $1.36E^{-5}$ | 0.1572 |
| C4 | *Rps3* | 0.44 | $2.42E^{-10}$ | $2.80E^{-6}$ |
| C4 | *Park7* | 2.24 | $7.82E^{-7}$ | 0.0090 |
| C4 | *Ms4a4c* | 0.63 | $6.03E^{-6}$ | 0.0697 |
| C5 | *Vim* | -0.75 | $7.76E^{-8}$ | 0.0009 |
| C5 | *Oaz1* | 0.38 | $6.24E^{-7}$ | 0.0072 |
| C5 | *Nt5c* | -1.10 | $5.59E^{-6}$ | 0.0646 |
| C5 | *Srpr* | -0.29 | $8.35E^{-6}$ | 0.0965 |
| C5 | *Rps9* | 0.42 | $1E^{-5}$ | 0.1252 |
| C6 | *Atp5c1* | -0.28 | $2.21E^{-9}$ | $2.55E^{-5}$ |
| C6 | *Trip11* | 2.98 | $4.76E^{-8}$ | 0.0005 |
| C6 | *Hint1* | -0.66 | $2.95E^{-7}$ | 0.0034 |
| C6 | *Rer1* | -0.95 | $7.15E^{-7}$ | 0.0083 |
| C6 | *Chmp6* | 1.79 | $3.14E^{-6}$ | 0.0363 |
| C6 | *Ifitm2* | 1.00 | $9.31E^{-6}$ | 0.1076 |
| C6 | *Serpinf1* | 2.91 | $1.25E^{-5}$ | 0.1444 |
| C6 | *X1110004f10rik* | 0.46 | $1.39E^{-5}$ | 0.1610 |

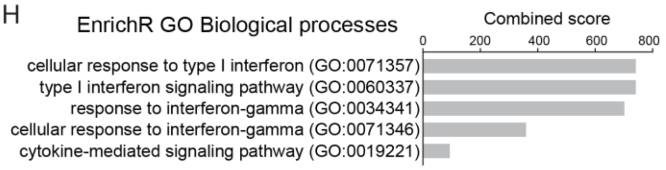


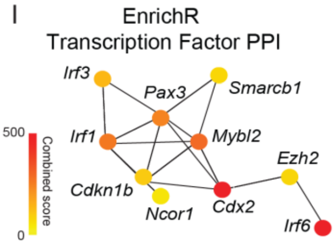

**Figure S6. IKKβ and IRF1 coordinate intra-tumoral cDC1 maturation.**

(**A**) Expression of selected genes related to DC maturation and cytokine expression on the UMAP space from Figure 6A.

(**B**) Jaccard indices showing the similarities between Seurat clusters of cDC1s from Figure 6A (purple) and Figure 2A (black).

(**C**) Violin plots showing the expression of *Ikbkb* and *Irf1* across Seurat clusters of Figure 6A.

(**D**) Scatter plots comparing fold change of DEGs in $Ikbkb^{\Delta Xcr1}$ (x axis) and $Irf1^{\Delta Xcr1}$ (y axis) in cDC1s from each cluster and cluster-independent DEGs. Coloured dots represent DEGs with adjusted *p* values <0.25. Spearman correlations between IKKβ and IRF1-dependent DEGs are shown.

(**E**) Violin plots showing the expression of selected DEGs related to antigen presentation, IFN response, DC maturation and TLR signaling in selected clusters of cDC1s from $Ikbkb^{\Delta Xcr1}$ and $Irf1^{\Delta Xcr1}$ mice.

(**F**) Expression of selected T cell chemokines across all cDC1 clusters (left) and differential expression in selected clusters from $Ikbkb^{\Delta Xcr1}$ and $Irf1^{\Delta Xcr1}$ mice (right).

(**G**) List of DEGs in $Ikbkb^{\Delta Xcr1}$ and $Irf1^{\Delta Xcr1}$ cDC1s with an adjusted *p* value <0.25 within specific clusters or independent of clusters (C-ind). Genes with a negative fold change are expressed to higher levels in WT cDC1s, and genes with a positive fold change are expressed to higher levels in mutant cDC1s.

(**H**) GO biological processes enriched in DEGs from scRNA-seq analysis of cDC1s from female $Ikbkb^{F/F}$ and $Ikbkb^{\Delta Xcr1}$ mice bearing YUMM1.7 tumors. Combined scores calculated by EnrichR analysis are shown.

(**I**) Enrichment of transcription factor PPI network in cDC1s from $Ikbkb^{F/F}$ versus $Ikbkb^{\Delta Xcr1}$ tumors based on scRNA-seq. The color intensity of each node is proportional to the combined score calculated by EnrichR.

### Supplementary Figure 7

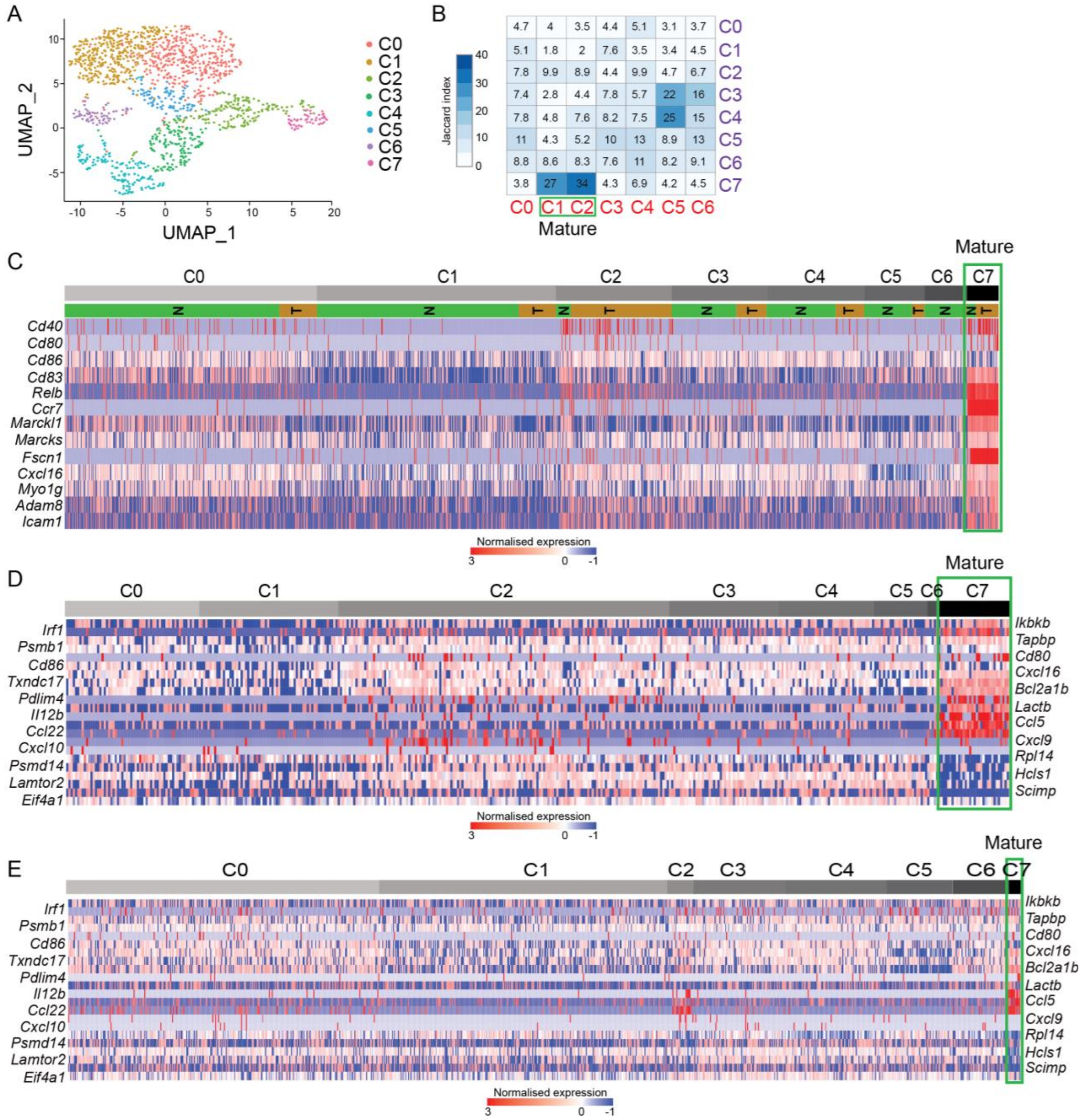

**Figure S7. Comparative analysis of cDC1 heterogeneity in lung adenocarcinoma and melanoma.**

(**A**) UMAP cell clustering using Seurat for 1418 cDC1s from naïve mouse lung tissue and lung adenocarcinoma.

(**B**) Jaccard indices showing the similarity between cDC1 clusters from mouse melanomas (red) and lung adenocarcinomas (purple).

(**C**) Gene expression heatmap of canonical cDC1 and DC maturation markers in clusters of cDC1s from naïve lungs (N) and adenocarcinomas (T).

(**D**,**E**) Gene expression heatmap of selected genes related to NF-κB and IRF1-dependent activation of cDC1s across clusters from (**D**) lung adenocarcinoma and (**E**) naïve lungs.
